# Neuronal activity promotes nuclear proteasome-mediated degradation of PDCD4 to regulate activity-dependent transcription

**DOI:** 10.1101/2021.02.06.429040

**Authors:** Wendy A. Herbst, Weixian Deng, James A. Wohlschlegel, Jennifer M. Achiro, Kelsey C. Martin

## Abstract

Activity-dependent gene expression is critical for synapse development and plasticity. To elucidate novel mechanisms linking neuronal activity to changes in transcription, we compared the nuclear proteomes of tetrodotoxin-silenced and bicuculline-stimulated cultured rodent neurons using nuclear-localized APEX2 proximity biotinylation and mass spectrometry. The tumor suppressor protein PDCD4 was enriched in the silenced nuclear proteome, and PDCD4 levels rapidly decreased in the nucleus and cytoplasm of stimulated neurons. The activity-dependent decrease of PDCD4 was prevented by inhibitors of both PKC and proteasome activity and by a phospho-incompetent mutation of Ser71 in the βTRCP ubiquitin ligase-binding motif of PDCD4. We compared the activity-dependent transcriptomes of neurons expressing wildtype or degradation-resistant (S71A) PDCD4. We identified 91 genes as PDCD4 targets at the transcriptional level, including genes encoding proteins critical for synapse formation, remodeling, and transmission. Our findings indicate that regulated degradation of nuclear PDCD4 facilitates activity-dependent transcription in neurons.

## Introduction

Stimulus-induced gene expression allows neurons to adapt their structure and function in response to a dynamically changing external environment (Gallegos et al., 2018; Holt et al., 2019; Yap & Greenberg, 2018). Activity-dependent transcription is critical to neural circuit function, from synapse formation during brain development (Flavell et al., 2006; Lin et al., 2008; Polleux et al., 2007; Wayman et al., 2006; West & Greenberg, 2011) to synaptic plasticity in the mature brain (Bloodgood et al., 2013; Chen et al., 2017; Ramanan et al., 2005; Tyssowski et al., 2018; Yap & Greenberg, 2018). Neuronal activity regulates gene expression at multiple levels, including chromatin modification and transcriptional regulation in the nucleus, as well as RNA localization, stability, and translation in the cytoplasm (Martin & Ephrussi, 2009). To produce activity-dependent changes in transcription, signals must be relayed from the site where the signal is received, at the synapse, to the nucleus. To better understand how neuronal activity is coupled with changes in transcription, we developed an assay to systematically identify activity-dependent changes in the nuclear proteome of neurons and therefore elucidate novel mechanisms by which neuronal activity alters the concentration of specific proteins in the nucleus.

Neuronal activity can change the concentration of nuclear proteins via a variety of mechanisms, from nucleocytoplasmic shuttling of signaling proteins, to synthesis and degradation of nuclear proteins (Bayraktar et al., 2020; Ch’ng et al., 2012; Dieterich et al., 2008; Lin et al., 2008; Ma et al., 2014; Upadhya et al., 2004). While the activity-dependent transcriptome and translatome of neurons has been characterized using RNA sequencing (Brigidi et al., 2019; Hrvatin et al., 2018; Lacar et al., 2016; Tyssowski et al., 2018) and TRAP-seq (Chen et al., 2017; Fernandez-Albert et al., 2019), little work has been done to characterize the population of proteins that undergo activity-dependent changes in nuclear abundance due to regulated transport or stability. By inhibiting translation to exclude changes due to protein synthesis, the present study focused on identifying pre-existing proteins that undergo activity-dependent changes in concentration in the nucleus via regulated nucleocytoplasmic trafficking and/or changes in stability.

Through our screen of nuclear proteins with activity-dependent changes in abundance, we discovered that **P**rogramme**d C**ell **D**eath **4**(PDCD4) undergoes a significant reduction in nuclear concentration following neuronal stimulation. PDCD4 has been studied primarily in the context of cancer, where it has been found to function as a tumor suppressor and translational inhibitor in the cytoplasm (Matsuhashi et al., 2019; Wang & Yang, 2018; Yang et al., 2003). These studies have revealed that the abundance of PDCD4 protein is regulated at multiple levels, including via translation (Asangani et al., 2008; Frankel et al., 2008; Ning et al., 2014), proteasome-mediated degradation (Dorrello et al., 2006), and nucleocytoplasmic trafficking (Böhm et al., 2003), with decreases in PDCD4 correlating with invasion, proliferation, and metastasis of many types of cancers (Allgayer, 2010; Chen et al., 2003; Wang & Yang, 2018; Wei et al., 2012).

Despite being expressed at significant levels in the brain, especially in the hippocampus and cortex (Lein et al., 2007; Li et al., 2020), few studies have addressed the role of PDCD4 in the nervous system. PDCD4 expression in neurons is altered by injury and stress (Jiang et al., 2017; Li et al., 2020; Narasimhan et al., 2013), and recent work has shown that, as in cancer cells, PDCD4 may act as a translational repressor in neurons (Di Paolo et al., 2020; Li et al., 2020). However, the impact of neuronal activity on PDCD4 concentration and the function of nuclear, as opposed to cytoplasmic PDCD4, remains unknown.

In this study, we describe an assay that represents the first, to our knowledge, to identify activity-dependent changes in the nuclear proteome of neurons, and does so in a manner that is independent of translation. Our results not only elucidate a novel mechanism by which activity can regulate the nuclear proteome, but they also support a role for the tumor suppressor protein PDCD4 in the nucleus during activity-dependent transcription in neurons.

## Results

### Identification of the nuclear proteome from silenced and stimulated neurons using APEX2 proximity biotinylation and mass spectrometry

To identify proteins that undergo activity-dependent changes in nuclear localization or abundance, we analyzed the nuclear proteomes of silenced and stimulated cultured rat forebrain neurons. In developing this assay, we used CREB Regulated Transcriptional Coactivator 1 (CRTC1) as a positive control, as we have previously shown that glutamatergic activity drives the synapse-to-nucleus import of CRTC1, and that neuronal silencing decreases CRTC1 nuclear abundance (Ch’ng et al., 2012, 2015). We initially used nuclear fractionation to capture the nuclear proteins but discovered that CRTC1 leaked out of the nucleus during the assay. This suggested that nuclear fractionation was not a suitable method and led us to instead use APEX2 proximity biotinylation (Hung et al., 2016), an *in situ* proximity ligation assay, to identify activity-dependent changes in the nuclear proteome. To specifically label the nuclear proteome, we fused the engineered ascorbate peroxidase APEX2 to two SV40 nuclear localization signals (NLSs, **Fig 1A**, Kalderon et al., 1984). APEX2 proximity ligation was advantageous for these experiments for the following reasons: 1) APEX2 biotinylated proteins can be captured directly by streptavidin pulldown, avoiding the need for subcellular fractionation, 2) biotinylation occurs rapidly (1-minute labeling period), and 3) APEX2 can be expressed in a specific cell type of interest. We designed a neuron-specific nuclear-localized APEX2 construct (**Fig 1A**) and transduced cultured rat forebrain neurons with adeno-associated virus (AAV) expressing APEX2-NLS. Immunofluorescence of transduced neurons revealed that APEX2-NLS was expressed specifically in the nucleus (**Fig 1B**). We optimized the multiplicity of infection of AAV to achieve high transduction efficiency without overexpression of the construct (important as higher doses of AAV led to APEX2-NLS expression in the cytoplasm). When all three components of the labeling reaction were supplied (APEX2-NLS, biotin-phenol, and H_2_O_2_), proteins were biotinylated specifically in neuronal nuclei (**Fig 1B**). No labeling was detected in the absence of APEX2-NLS, biotin-phenol, or H_2_O_2_.

**Figure 1.**
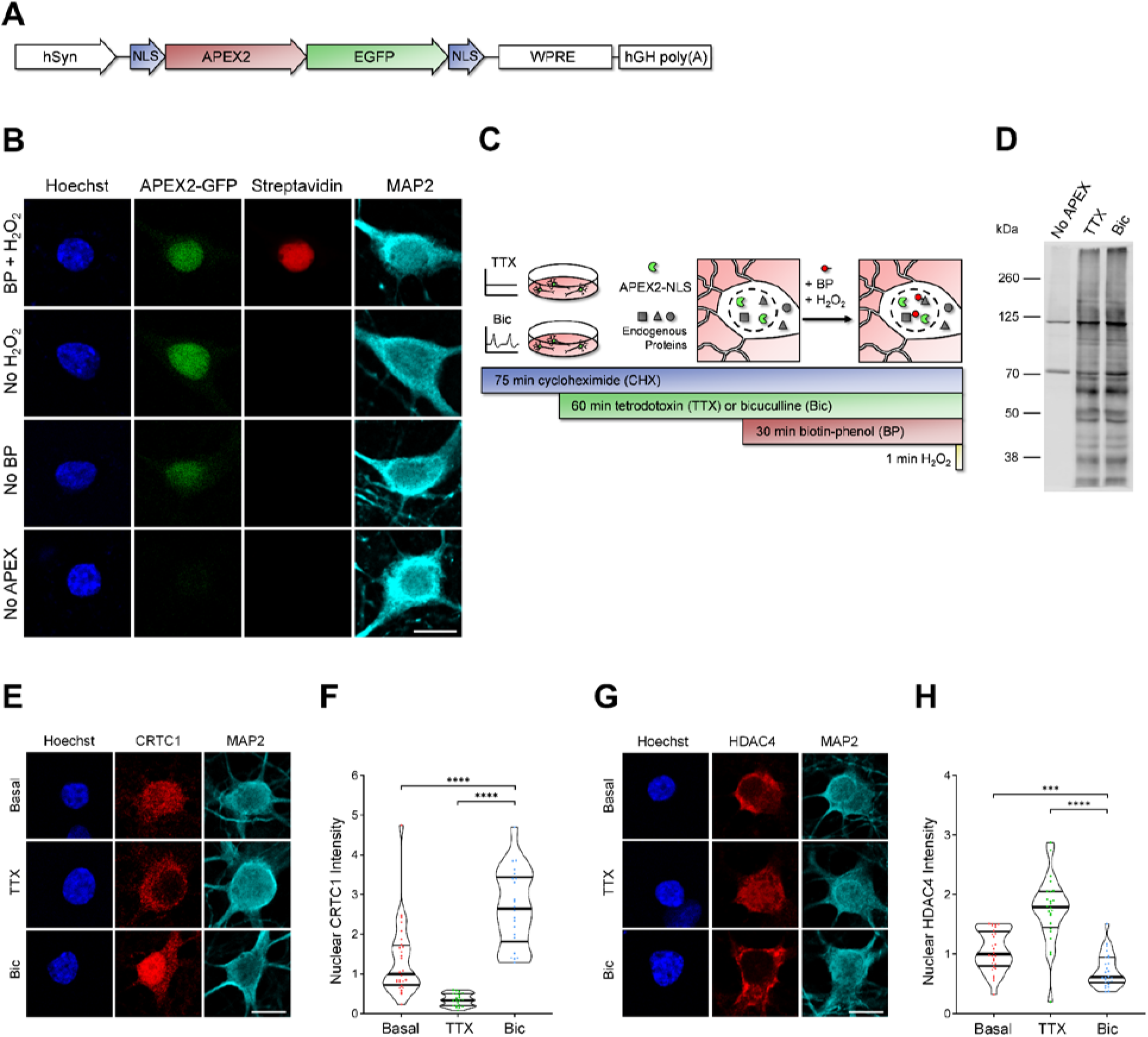
Identification of the nuclear proteomes from silenced and stimulated neurons using APEX2 proximity biotinylation. **A)** Design of APEX2-NLS construct. hSyn = human synapsin promoter, NLS = SV40 nuclear localization signal. **B)** Immunocytochemistry (ICC) of cultured rat forebrain neurons after APEX2 proximity biotinylation labeling. Nuclear proteins were biotinylated (streptavidin, red) by the combined presence of APEX2-NLS (GFP, green), biotin-phenol (BP), and H_2_O_2_. Scale bar = 10 μm. **C)** Workflow for labeling nuclear proteins from silenced and stimulated neurons. APEX2 labeling diagram based on (Hung et al., 2016). **D)** Western blot of cultured neuron protein lysates from No APEX, APEX+TTX, or APEX+Bic conditions, stained with streptavidin. **E)** CRTC1 ICC of basal, TTX-silenced, and Bic-stimulated neurons. Scale bar = 10 μm. **F)** Violin plots of normalized nuclear CRTC1 ICC intensity. Basal n = 28, TTX n = 21, Bic n = 22 cells, from 1 set of cultures. **G)** HDAC4 ICC of basal, TTX-silenced, and Bic-stimulated neurons. Scale bar = 10 μm. **H)** Violin plots of normalized nuclear HDAC4 ICC intensity. Basal n = 28, TTX n = 24, Bic n = 26 cells, from 1 set of cultures. Statistical significance is indicated by ***p < 0.001 and ****p < 0.0001, from Mann-Whitney U test with Bonferroni correction.

To identify proteins that undergo activity-dependent changes in nuclear abundance, we silenced neurons for 1 hour with the voltage-gated sodium channel antagonist tetrodotoxin (TTX) or stimulated neurons for 1 hour with bicuculline (Bic), which inhibits GABA_A_ receptors and drives glutamatergic transmission. We also inhibited protein synthesis using cycloheximide (CHX) in these experiments because many of the genes that are rapidly transcribed and translated in response to activity encode nuclear proteins (Alberini, 2009; D. A. Heinz & Bloodgood, 2020; Yap & Greenberg, 2018). We were concerned that the translation of activity-dependent genes would overshadow – and thereby hinder the detection of – activity-dependent changes in the nuclear proteome resulting from alterations in nuclear protein localization or stability.

After silencing or stimulating the neurons for 1 hour and performing the 1-minute labeling reaction (**Fig 1C)**, protein lysates were collected for analysis by western blot and mass spectrometry. In neurons expressing APEX2-NLS, many proteins at a variety of molecular weights were biotinylated in both TTX and Bic conditions, while very few proteins were biotinylated in the No-APEX control, as detected by western blot (**Fig 1D**). The bands detected in the No-APEX control were at molecular weights of known endogenously biotinylated proteins (Hung et al., 2016). For mass spectrometry, biotinylated proteins were captured using streptavidin pulldown and the nuclear proteomes were characterized using the TMT-MS3-SPS acquisition method (Ting et al., 2011) through LC-MS. We detected 4,407 proteins, and of those, 2,860 proteins were significantly enriched above the No-APEX negative control with log2 fold change (FC) > 3 and adjusted p-value < 0.05 (**Table S1**). When comparing the Bic and TTX conditions, 23 proteins were differentially expressed in the nucleus with log2FC > 0.5 (for Bic) or log2FC < −0.5 (for TTX) with p-value < 0.05 (**Table S1**). The highest-ranked protein by log2FC enriched in the Bic versus TTX nuclear proteome was the synapse-to-nucleus signaling protein, CRTC1 (Ch’ng et al., 2012, 2015; Nonaka et al., 2014; Sekeres et al., 2012). The highest-ranked proteins enriched in the TTX versus Bic nuclear proteome were HDAC4 and HDAC5, both of which have been reported to undergo nuclear export following neuronal stimulation (Chawla et al., 2003; Schlumm et al., 2013). Using immunocytochemistry (ICC), we confirmed that CRTC1 increased in the nucleus (median normalized intensity Basal = 1.00, TTX = 0.33, Bic = 2.64; Basal vs Bic p < 0.0001, TTX vs Bic p < 0.0001) and HDAC4 decreased in the nucleus (median normalized intensity Basal = 1.00, TTX = 1.79, Bic = 0.61; Basal vs Bic p = 0.0006, TTX vs Bic p < 0.0001) following Bic stimulation (**Fig 1E-H**).

### Neuronal stimulation decreases PDCD4 protein concentration in the nucleus and cytoplasm of neurons

Among the proteins that underwent activity-dependent changes in nuclear concentration, the PDCD4 protein was significantly enriched in the TTX-treated nuclear proteome compared to the Bic-treated nuclear proteome. To validate this finding, we characterized the expression of PDCD4 protein in cultured neurons using ICC, and found that PDCD4 was present both in the nucleus and cytoplasm of neurons (**Fig 2A, Fig S1A**). Bic stimulation significantly decreased PDCD4 protein expression in the nucleus by ~50% (median normalized intensity Basal = 1.00, TTX = 1.00, Bic = 0.55; Basal vs Bic p < 0.0001, TTX vs Bic p < 0.0001; **Fig 2B**), with a smaller decrease of ~20% in the cytoplasm (median normalized intensity Basal = 1.00, TTX = 1.07, Bic = 0.81; Basal vs Bic p < 0.0001, TTX vs Bic p < 0.0001; **Fig 2C**). The decrease of PDCD4 occurred within 15 minutes of Bic stimulation, and PDCD4 protein levels continued to decrease with longer incubations of Bic (**Fig S1B**). After washout of a 1-hour Bic stimulation, PDCD4 protein expression gradually returned to baseline levels, although only half of the PDCD4 protein concentration was restored at 24 hours (**Fig S1C**).

**Figure 2.**
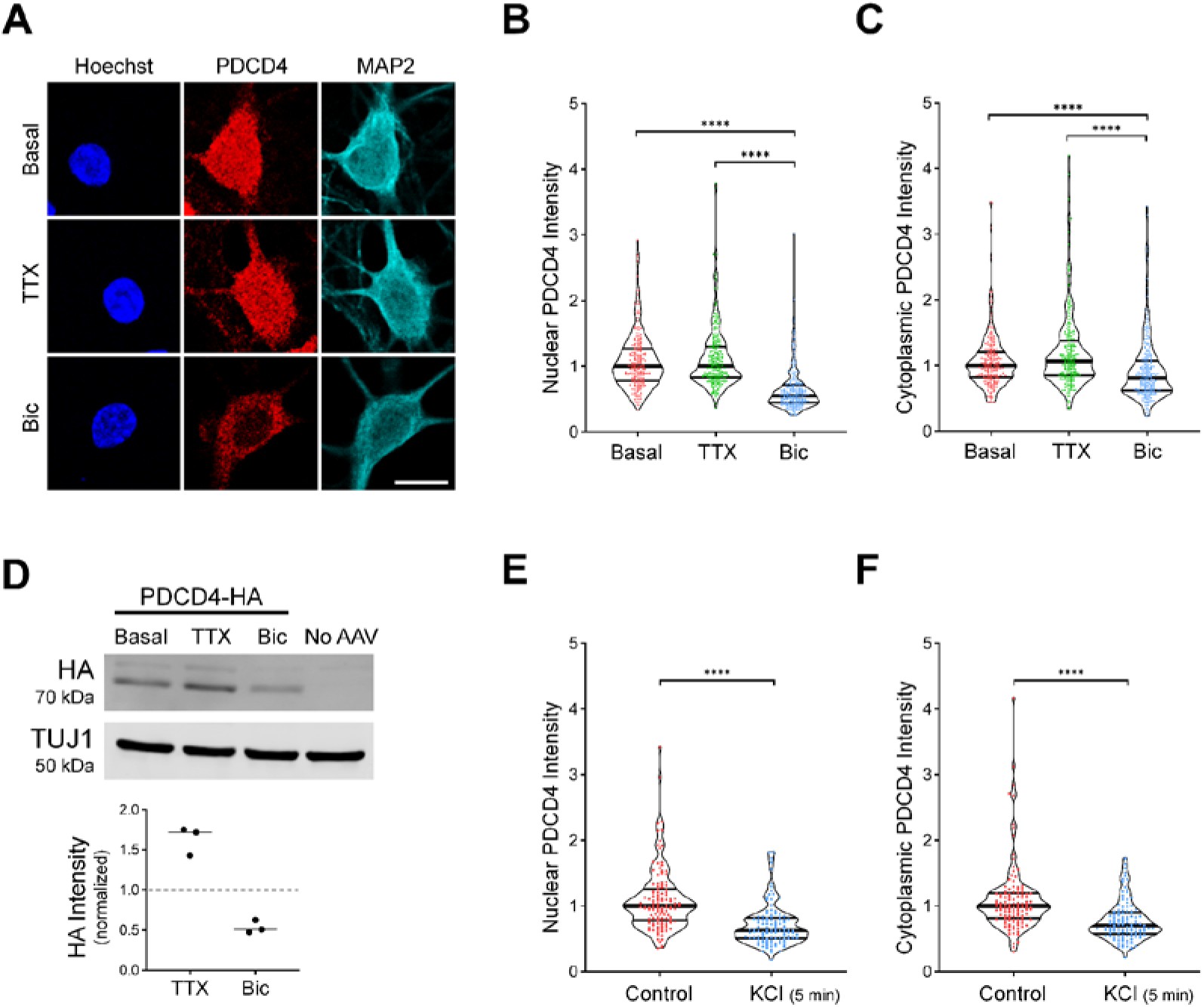
Neuronal stimulation decreases PDCD4 protein concentration in the nucleus and cytoplasm of neurons. **A)** PDCD4 immunocytochemistry (ICC) of basal, TTX-silenced, and Bic-stimulated neurons. Scale bar = 10 μm. **B)** Violin plots of normalized nuclear PDCD4 ICC intensity. Basal n = 226, TTX n = 227, Bic n = 218 cells, from 6 sets of cultures. **C)** Violin plots of normalized cytoplasmic PDCD4 ICC intensity in the same cells as in B. **D)** Top: Western blot of protein lysates from basal, TTX-silenced, and Bic-stimulated neurons transduced with PDCD4-HA AAV. Bottom: Quantification of western blot, from 3 sets of cultures. HA intensity was normalized to TUJ1 intensity. Within each experiment, all samples were normalized to the basal sample. **E)** Violin plots of normalized nuclear PDCD4 ICC intensity. Control n = 118, KCl n = 110 cells, from 3 sets of cultures. **F)** Violin plots of normalized cytoplasmic PDCD4 ICC intensity in the same cells as in G. Statistical significance is indicated by **p < 0.01, ***p < 0.001, and ****p < 0.0001, from Mann-Whitney U test with Bonferroni correction.

In complementary experiments, we transduced neurons with AAV expressing C-terminal HA-tagged PDCD4 (**Fig S1D**) and characterized PDCD4-HA expression by western blot (**Fig 2D**) and ICC (**Fig S1E-F**). By western blot, we found that total PDCD4-HA protein levels decreased by ~50% following Bic stimulation (median normalized intensity TTX/Basal = 1.72, Bic/Basal = 0.48; **Fig 2D**). By ICC, both nuclear and cytoplasmic PDCD4-HA decreased by ~40% (**Fig S1E-F**). To further validate the Bic-induced decrease in PDCD4, we also created an N-terminal V5-tagged PDCD4 plasmid (**Fig S1D**) and transfected the construct in neurons. Consistent with the results from endogenous PDCD4 and transduced C-terminally HA-tagged PDCD4, both nuclear and cytoplasmic V5-PDCD4 decreased by ~40% with Bic stimulation (**Fig S1G-H**).

We also found that depolarization of neurons with 40 mM KCl for 5 min significantly decreased PDCD4 protein concentration by ~40% in the nucleus (median normalized intensity Control = 1.00, KCl = 0.63; Control vs KCl p < 0.0001; **Fig 2E**) and ~30% in the cytoplasm (median normalized intensity Control = 1.00, KCl = 0.70; Control vs KCl p < 0.0001; **Fig 2F**), as detected immediately after the 5 min treatment. These results indicate that increases in glutamatergic transmission and depolarization lead to a rapid and long-lasting reduction in PDCD4 abundance in the nucleus of neurons.

### PDCD4 undergoes proteasome-mediated degradation following neuronal stimulation

In non-neuronal cells, PDCD4 has been reported to undergo miRNA-mediated translational repression (Asangani et al., 2008; Frankel et al., 2008; Ning et al., 2014), stimulus-induced nuclear export (Böhm et al., 2003), and stimulus-induced proteasome-mediated degradation (Dorrello et al., 2006). Since the assay we used to detect activity-dependent changes in the nuclear proteome was conducted in the presence of the protein synthesis inhibitor CHX, we considered it unlikely that the Bic-induced decrease in PDCD4 was due to miRNA-mediated translational repression. To confirm this, we conducted PDCD4 ICC of TTX-silenced and Bic-stimulated neurons in the presence or absence of CHX (**Fig S2A-B**). While CHX potently blocked the activity-dependent increase in FOS immunoreactivity (**Fig S2C**), it did not block the Bic-induced decrease in PDCD4 (**Fig S2A-B**), thereby ruling out a role for activity-dependent miRNA-mediated translational regulation.

To investigate the mechanism underlying the decrease in nuclear PDCD4, we tested if the decrease in nuclear abundance was due to activity-dependent increases in nuclear export. Nuclear export of PDCD4 is mediated by the nuclear export protein, CRM1, and sensitive to the nuclear export inhibitor, leptomycin B (LMB) (Böhm et al., 2003). Long incubation with LMB successfully caused nuclear accumulation of PDCD4 in unstimulated neurons (**Fig S2D**), and yet LMB was unable to prevent the activity-dependent decrease of nuclear PDCD4 following stimulation (median normalized intensity Basal = 1.00, TTX = 1.08, Bic = 0.59, LMB-Basal = 1.30, LMB-TTX = 1.16, LMB-Bic = 0.52; Basal vs Bic p < 0.0001, TTX vs Bic p < 0.0001, LMB-Basal vs LMB-Bic p < 0.0001, LMB-TTX vs LMB-Bic p < 0.0001; **Fig 3A**, **Fig S2E**). This result demonstrates that regulated nuclear export is not required for the activity-dependent decrease of PDCD4.

**Figure 3.**
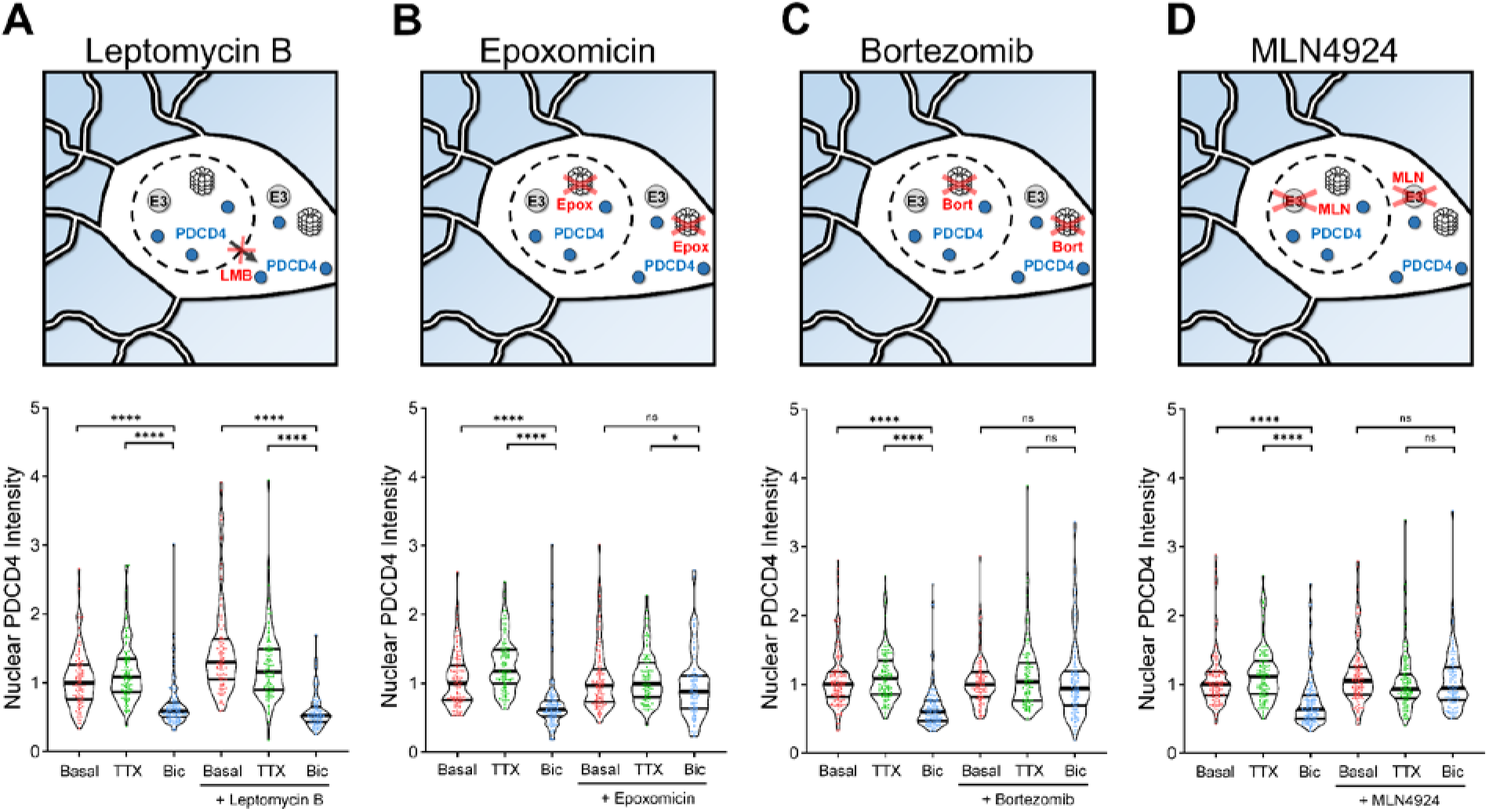
PDCD4 undergoes proteasome-mediated degradation –not nuclear export– following neuronal stimulation. **A)** Top: Schematic of CRM1-mediated nuclear export inhibitor leptomycin B (LMB) experiments. Bottom: Violin plots of normalized nuclear PDCD4 immunocytochemistry (ICC) intensity. Basal n = 109, TTX n = 109, Bic n = 109, LMB-Basal n = 106, LMB-TTX n = 100, LMB-Bic n = 86 cells, from 3 sets of cultures. **B)** Top: Schematic of proteasome inhibitor epoxomicin (epox) experiments. Bottom: Violin plots of normalized nuclear PDCD4 ICC intensity. Basal n = 113, TTX n = 106, Bic n = 103, Epox-Basal n = 107, Epox-TTX n = 94, Epox-Bic n = 86 cells, from 3 sets of cultures. **C)** Top: Schematic of proteasome inhibitor bortezomib (bort) experiments. Bottom: Violin plots of normalized nuclear PDCD4 ICC intensity. Basal n = 131, TTX n = 111, Bic n = 120, Bort-Basal n = 100, Bort-TTX n = 98, Bort-Bic n = 116 cells, from 3 sets of cultures. **D)** Top: Schematic of MLN4924 (MLN) experiments. MLN4924 inhibits neddylation, preventing the activation of Cullin-RING E3 ubiquitin ligases. Bottom: Violin plots of normalized nuclear PDCD4 ICC intensity. Basal n = 130, TTX n = 120, Bic n = 120, MLN-Basal n = 115, MLN-TTX n = 108, MLN-Bic n = 97 cells, from 3 sets of cultures. Statistical significance is indicated by *p < 0.05 and ****p < 0.0001, from Mann-Whitney U test with Bonferroni correction.

We hypothesized that regulated ubiquitin proteasome-mediated degradation may explain the Bic-induced decrease of PDCD4. To test this idea, we incubated TTX and Bic-stimulated neurons with the proteasome inhibitors epoximicin (Epox) or bortezomib (Bort). As shown in **Fig 3B-C** and **Fig S2F-G**, the proteasome inhibitors impaired Bic-induced decreases of PDCD4 in both nucleus and cytoplasm of neurons, indicating that neuronal activity decreases PDCD4 concentrations via proteasome-mediated degradation (**Fig 3B:** median normalized intensity Basal = 1.00, TTX = 1.18, Bic = 0.62, Epox-Basal = 0.97, Epox-TTX = 0.99, Epox-Bic= 0.88; Basal vs Bic p < 0.0001, TTX vs Bic p < 0.0001, Epox-Basal vs Epox-Bic p = 0.1662, Epox-TTX vs Epox-Bic p = 0.0174. **Fig 3C:** median normalized intensity Basal = 1.00, TTX = 1.09, Bic = 0.60, Bort-Basal = 1.00, Bort-TTX = 1.04, Bort-Bic = 0.94; Basal vs Bic p < 0.0001, TTX vs Bic p < 0.0001, Bort-Basal vs Bort-Bic p = 0.6224, Bort-TTX vs Bort-Bic p = 0.2648). The finding that the nuclear export inhibitor LMB did not block the Bic-induced decrease of PDCD4 in the nucleus (**Fig 3A**) indicates that activity regulates proteasome-mediated degradation of nuclear PDCD4 directly in the nucleus, rather than by nuclear export of PDCD4 followed by degradation in the cytoplasm.

The E3 ubiquitin ligases βTRCP1/2 have been shown to be required for the proteasome-mediated degradation of PDCD4 in the T98G glioblastoma cell line (Dorrello et al., 2006). βTRCP1/2 belong to the family of Cullin-RING E3 ubiquitin ligases, which require neddylation in order to be activated (Merlet et al., 2009). To determine if this family of ligases is involved in the activity-dependent decrease of PDCD4 in neurons, we used the neddylation inhibitor MLN4924 (MLN) and found that it blocked the Bic-induced decrease of PDCD4 in the nucleus (**Fig 3D**) and cytoplasm (**Fig S2H**) of neurons (**Fig 3D**: median normalized intensity Basal = 1.00, TTX = 1.11, Bic = 0.64, MLN-Basal = 1.05, MLN-TTX = 0.93, MLN-Bic = 0.95; Basal vs Bic p < 0.0001, TTX vs Bic p < 0.0001, MLN-Basal vs MLN-Bic p = 0.329, MLN-TTX vs MLN-Bic p = 1). This result further supports the finding that PDCD4 undergoes proteasome-mediated degradation following Bic stimulation, likely through ubiquitination by βTRCP1/2.

### PDCD4 S71A mutation and PKC inhibition prevent the activity-dependent decrease of PDCD4

PDCD4 contains a canonical βTRCP-binding motif, and a single phospho-incompetent serine-to-alanine mutation at either Ser67, Ser71, or Ser76 has been shown to prevent the stimulus-induced degradation of PDCD4 in T98G glioblastoma cells (Dorrello et al., 2006). Dorrello et al. demonstrated that PDCD4 must be phosphorylated at these sites in order to interact with βTRCP and undergo subsequent proteasome-mediated degradation. To test if a mutation in PDCD4 at one of these sites would prevent the activity-dependent decrease of PDCD4 in neurons, we expressed wild-type (WT) and mutant (S71A) PDCD4-HA in cultured neurons using AAV (**Fig 4A-B**). Similar to endogenous PDCD4, we found that WT PDCD4-HA decreased following Bic stimulation, with a ~30% decrease in nuclear HA intensity and a ~15% decrease in cytoplasmic intensity, as detected by ICC (**Fig 4C**, **Fig S3A**). In contrast, the PDCD4-HA S71A mutant did not undergo an activity-dependent decrease in either the nucleus or cytoplasm, but showed a slight activity-dependent increase in nuclear intensity (**Fig 4C**: median normalized intensity WT-Basal = 1.00, WT-TTX = 1.13, WT-Bic = 0.67, S71A-Basal = 1.05, S71A-TTX = 1.21, S71A-Bic = 1.27; WT-Basal vs WT-Bic p < 0.0001, WT-TTX vs WT-Bic p < 0.0001, S71A-Basal vs S71A-Bic p = 0.0034, S71A-TTX vs S71A-Bic p = 0.14). In complementary experiments, we found that Bic stimulation resulted in a ~40% decrease of WT PDCD4-HA signal by western blot, while PDCD4-HA S71A did not change after stimulation (**Fig 4D**: median normalized intensity WT Bic/Basal = 0.59, S71A Bic/Basal = 0.90).

**Figure 4.**
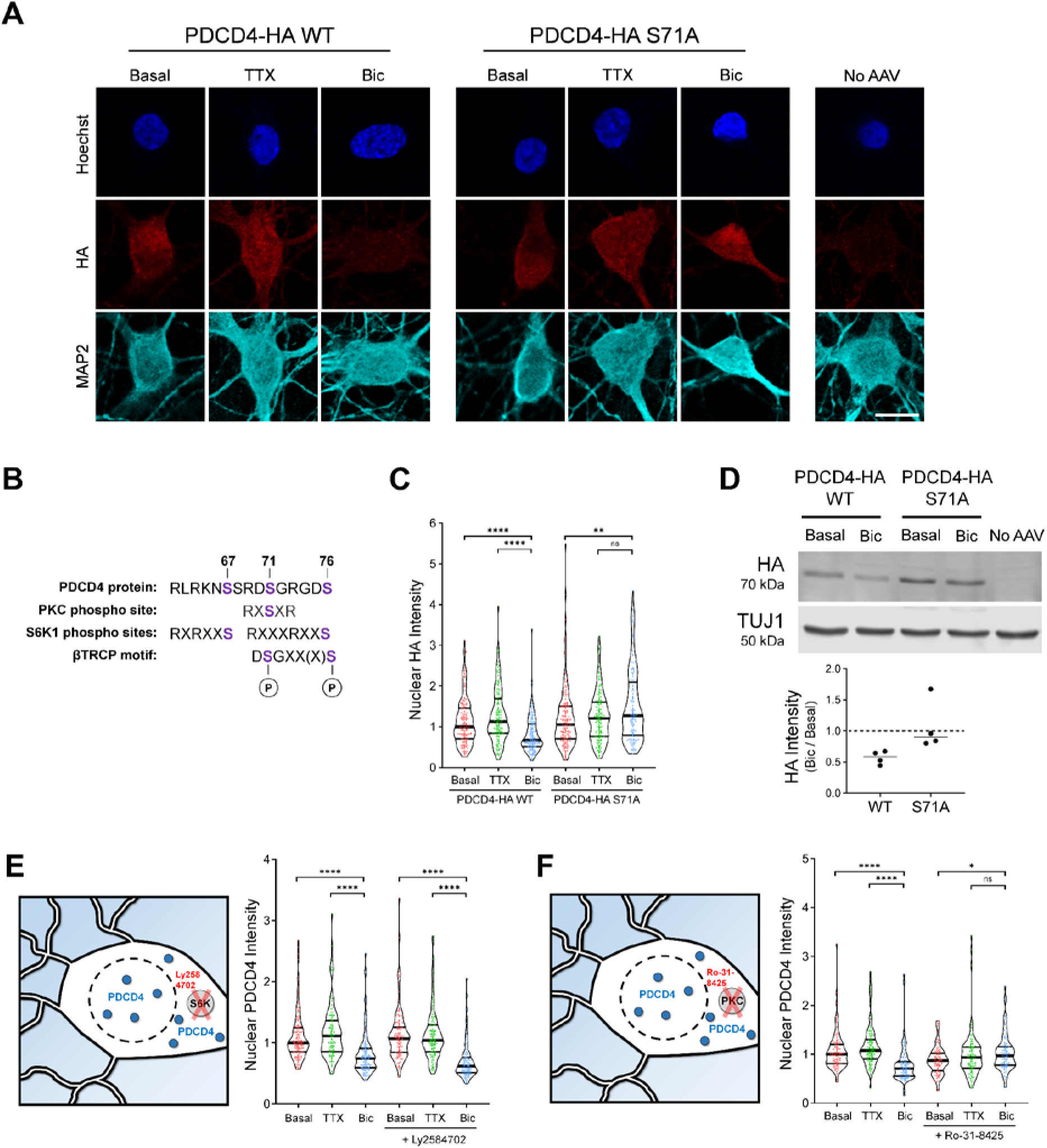
PDCD4 S71A mutation and PKC inhibition prevent the activity-dependent decrease of PDCD4. **A)** HA immunocytochemistry (ICC) of basal, TTX-silenced, and Bic-stimulated neurons transduced with PDCD4-HA WT, PDCD4-HA S71A AAV, or No AAV (negative control). Scale bar = 10 μm. **B)** PDCD4 protein sequence (amino acids 62-76). PKC and S6K1 phosphorylation sites are indicated in purple. Adapted from (Matsuhashi et al., 2019). **C)** Violin plots of normalized nuclear HA ICC intensity. WT-Basal n = 144, WT-TTX n = 136, WT-Bic n = 147, S71A-Basal n = 158, S71A-TTX n = 140, S71A-Bic n = 122 cells, from 4 sets of cultures. **D)** Top: Western blot of protein lysates from basal and Bic-stimulated neurons transduced with PDCD4-HA WT or PDCD4-HA S71A. Bottom: Quantification of western blot, from 4 sets of cultures. HA intensity was normalized to TUJ1 intensity. Within each experiment, each Bic sample was normalized to its respective basal sample. **E)** Left: Schematic of S6K inhibitor Ly2584702 (LY) experiments. Right: Violin plots of normalized nuclear PDCD4 ICC intensity. Basal n = 138, TTX n = 104, Bic n = 122, LY-Basal n = 112, LY-TTX n = 112, LY-Bic n = 107 cells, from 3 sets of cultures. **F)** Left: Schematic of PKC inhibitor Ro-31-8425 (Ro) experiments. Right: Violin plots of normalized nuclear PDCD4 ICC intensity. Basal n = 101, TTX n = 101, Bic n = 96, Ro-Basal n = 81, Ro-TTX n = 95, Ro-Bic n = 87 cells, from 3 sets of cultures. Statistical significance is indicated by *p < 0.05, **p < 0.01, and ****p < 0.0001, from Mann-Whitney U test with Bonferroni correction.

The βTRCP-binding motif is adjacent to a canonical phosphorylation consensus site for the kinase S6K1 (**Fig 4B**), and Dorrello et al. demonstrated that knockdown of S6K1 prevented the stimulus-induced degradation of PDCD4 in glioblastoma cells. In neurons, however, we found that the S6K inhibitor LY2584702 had no effect on the Bic-induced decrease of PDCD4 (**Fig 4E**, **Fig S3B**. **Fig 4E**: median normalized intensity Basal = 1.00, TTX = 1.11, Bic = 0.74, LY-Basal = 1.07, LY-TTX = 1.04, LY-Bic = 0.62; Basal vs Bic p < 0.0001, TTX vs Bic p < 0.0001, LY-Basal vs LY-Bic p < 0.0001, LY-TTX vs LY-Bic p < 0.0001), despite its ability to potently inhibit the phosphorylation of a known S6K target, ribosomal protein S6 (**Fig S3C**). In Huh7 hepatoma cells, mTOR/S6K and phospho-Ser67 are required for epidermal growth factor (EGF)-induced degradation of PDCD4, whereas protein kinase C (PKC) and phospho-Ser71 are required for 12-O-tetradecanoylphorbol-13-acetate (TPA)-induced degradation of PDCD4 (Matsuhashi et al., 2014, 2019; Nakashima et al., 2010). Of note, Ser71 is a consensus site for PKC, while Ser67 and Ser76 are consensus sites for S6K1 (**Fig 4B**) (Dorrello et al., 2006; Matsuhashi et al., 2019). We thus hypothesized that PKC may be important for the activity-dependent degradation of PDCD4 through phosphorylation of Ser71. To test this idea, we used the pan-PKC inhibitor Ro-31-8425 and found that it completely prevented the activity-dependent decrease (and slightly increased nuclear intensity) of PDCD4 (**Fig 4F**, **Fig S3D, Fig 4F**: median intensity Basal = 1.00, TTX = 1.08, Bic = 0.71, Ro-Basal = 0.87, Ro-TTX = 0.93, Ro-Bic = 0.97; Basal vs Bic p < 0.0001, TTX vs Bic p < 0.0001, Ro-Basal vs Ro-Bic p = 0.0114, Ro-TTX vs Ro-Bic p = 0.3892). These results suggest that in response to neuronal activity, PKC phosphorylates PDCD4 at Ser71, which enables the ubiquitin ligase βTRCP1/2 to bind and promote the proteasome-mediated degradation of PDCD4 in the nucleus.

### Stimulus-induced degradation of PDCD4 regulates the expression of neuronal activity-dependent genes

The finding that PDCD4 protein concentration is dynamically regulated in the nucleus in response to synaptic activity points to a possible nuclear function for PDCD4. To investigate a role for PDCD4 in the regulation of activity-dependent transcription, we performed RNA-seq of forebrain cultures transduced with either wild-type PDCD4 or degradation-resistant PDCD4 (S71A), following neuronal silencing with TTX or stimulation with Bic for 1 hour (**Fig 5; Table S2**). Given that PDCD4 has a well-known role in regulating translation, we sought to distinguish between direct PDCD4-mediated transcriptional changes in the nucleus and the changes in expression that are downstream of PDCD4-mediated translational changes in the cytoplasm by performing the experiments in either the presence or absence of CHX. We detected robust Bic-induced increases in normalized read counts of transcripts for canonical immediate early genes such as *Npas4*, *Rgs2*, and *Egr4* in all biological replicates (**Fig 5A**). We identified 912 activity-dependent genes, defined as genes with significant differential expression between Bic and TTX for PDCD4 WT samples (459 upregulated, 453 downregulated; adjusted p-value < 0.05 for WT no CHX; **Fig 5B-D**). Clustering of activity-dependent genes by fold change across sample type revealed that most activity-dependent genes showed similar fold changes between PDCD4 WT and PDCD4 S71A samples (**Fig 5B**), especially for genes with relatively high activity-dependent fold changes (**Fig 5C**). However, **Fig 5B** also shows that PDCD4 S71A altered activity-dependent changes in gene expression for a subset of genes. Specifically, we found that PDCD4 S71A led to a decrease in activity-induced differential expression for a substantial proportion of genes: 43% of activity-dependent upregulated genes (198 genes) and 57% of activity-dependent downregulated genes (260 genes) were not significantly upregulated or downregulated, respectively, in the PDCD4 S71A samples (**Fig 5D**). These results suggest that regulated degradation of PDCD4 is important for the expression of activity-dependent genes in neurons.

**Figure 5.**
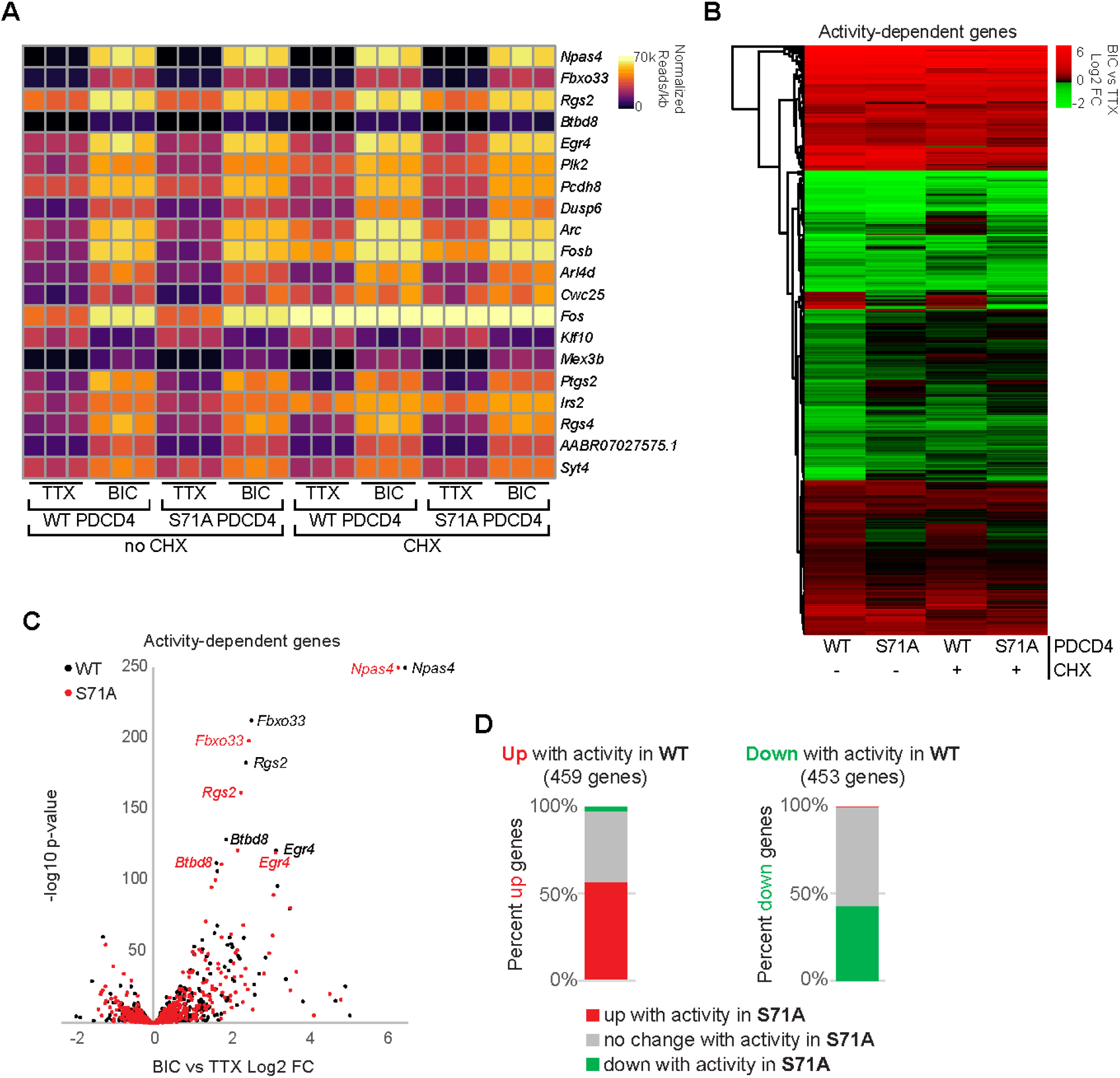
Stimulus-induced degradation of PDCD4 regulates the expression of neuronal activity-dependent genes. **A)** For each biological replicate, normalized read counts (from DESeq2) divided by transcript length are shown for the top 20 activity-dependent genes, ranked by adjusted p-value for PDCD4 WT no CHX samples. Each row represents a gene, and each column represents a biological replicate. The color of each box indicates transcript abundance (note: color is not scaled linearly in order to display full range of read counts; see **Table S2** for full dataset). **B)** Stimulation-induced log2 fold change (FC) for all 912 activity-dependent genes, clustered by fold change across sample type. Each row represents a gene and each column represents a sample type. The color legend represents Bic vs TTX log2FC with red representing upregulation, green representing downregulation, and black indicating log2FC of zero. **C)** For each activity-dependent gene, Bic versus TTX log2FC is plotted against −log10 of adjusted p-value, from PDCD4 WT samples (black) and PDCD4 S71A samples (red). Gene names for the top five activity-dependent genes by adjusted p-value are labeled. For both PDCD4 WT and PDCD4 S71A samples, *Npas4* Bic vs TTX adjusted p-value was zero (-log10 of zero is not defined), therefore for display, the −log10 adjusted p-value for *Npas4* was set to 250 for both samples. **D)** Activity-dependent upregulated genes (left bar) and activity-dependent downregulated genes (right bar) were categorized by the activity-dependent differential expression in PDCD4 S71A samples. The colors in each bar show the percentage of activity-dependent genes showing activity-dependent upregulation (red), no change (gray) or downregulation (green) in PDCD4 S71A samples.

This inhibition of activity-dependent gene expression could be due to both a potential role for PDCD4 in transcriptional processes and secondary effects from PDCD4’s regulation of translation of specific transcripts (Matsuhashi et al., 2019; Wang & Yang, 2018). To isolate effects at the transcriptional level, we focused on CHX-insensitive activity-dependent genes, that is, genes that showed activity-dependent differential expression in both the presence and absence of CHX (in WT, 459 genes after excluding 3 genes that showed differential expression in different directions +/− CHX). We ranked CHX-insensitive genes by their change in activity-dependent fold change between PDCD4 WT and PDCD4 S71A samples and identified 91 putative PDCD4 target genes that showed large differences in activity-dependent gene expression between wild-type and degradation-resistant PDCD4 samples (see Methods; **Fig 6A**). We validated the effect of PDCD4 on activity-dependent gene expression with RT-qPCR for two of the genes with the largest change between PDCD4 WT and PDCD4 S71A (*Scd1* and *Thrsp*; **Fig S4A**). We performed motif analysis of promoter sequences of the putative PDCD4 target genes and found similar motifs as in promoters of other CHX-insensitive activity-dependent genes (e.g. AP-1/TRE, ATF/CRE, Sp1/Klf motifs; **Fig S4B**), suggesting there was not a specific transcription factor motif associated with putative PDCD4 targets genes. Gene ontology (GO) analysis of the putative PDCD4 target genes showed enrichment for neuronal signaling terms such as “nervous system development” (GO:0007399; 28 genes; FDR = 9.28E-04) and “synapse” (GO:0045202; 18 genes; FDR = 6.54E-03), whereas, for comparison, other CHX-insensitive activity-dependent genes showed enrichment for terms related to transcription such as “regulation of gene expression” (GO:0010468; 169 genes; FDR = 3.11E-25) and “nuclear chromosome” (GO:0000228; 51 genes; FDR = 2.88E-09; **Fig 6B**, **Table S3**). Putative PDCD4 targets included genes encoding proteins critical for synapse formation, remodeling and transmission such as Shank1, p35, Abhd17b, Gap43, Cofilin, Spectrin-β2, Myosin-Va, Dendrin, Jacob, SNAP-β, Voltage-dependent calcium channel-α2/δ1, α-tubulin, and β-actin (**Fig 6C**). Together, these results suggest that PDCD4 functions in the nucleus to regulate the expression of a subset of genes, and that inhibiting the stimulation-induced degradation of PDCD4 results in a suppression of the transcription of many activity-dependent genes important for neuronal synaptic function (**Fig S5**).

**Figure 6.**
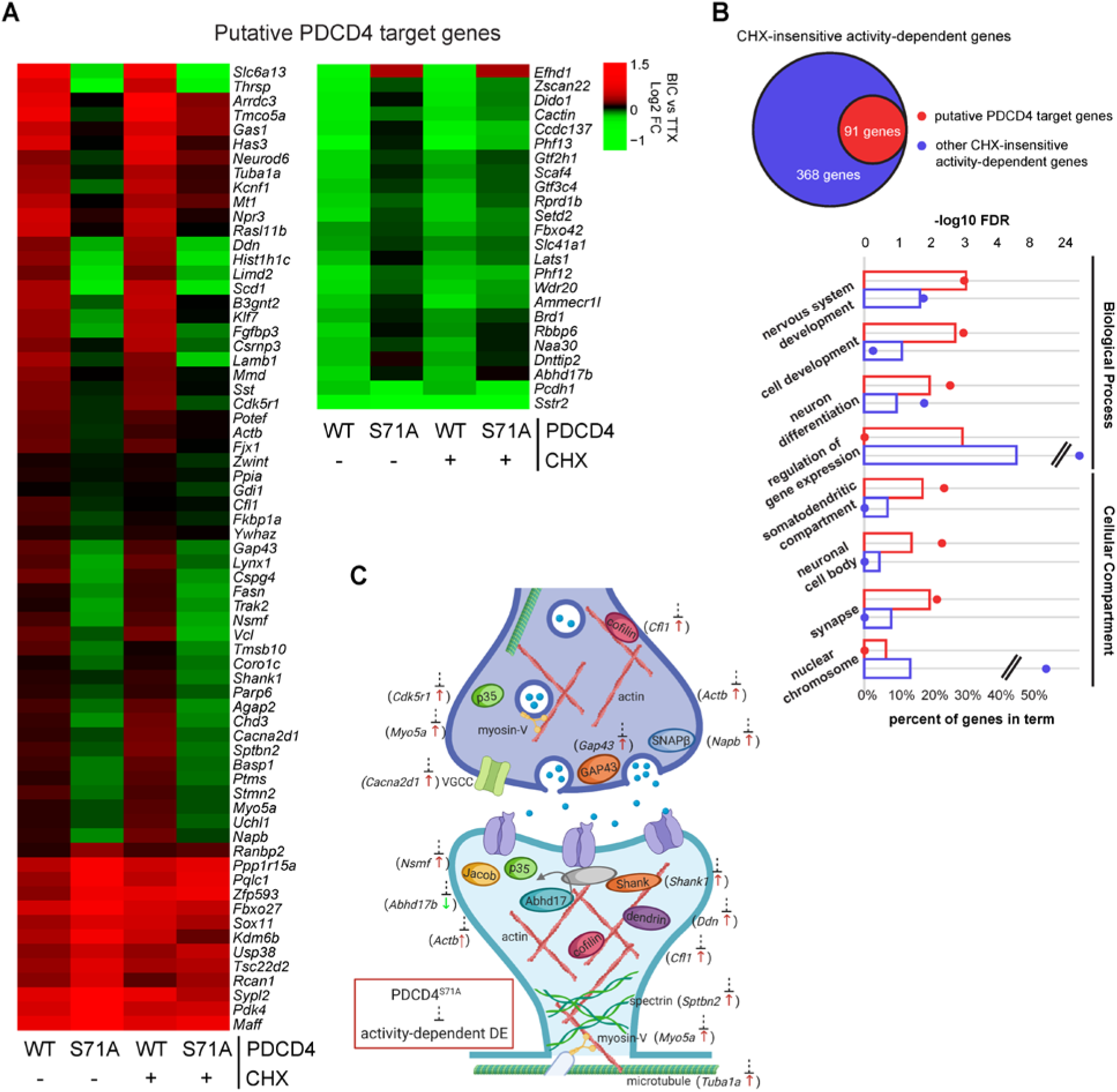
Degradation-resistant PDCD4 suppresses activity-dependent changes in expression of synaptic genes. **A)** Bic versus TTX log2FC for all 91 putative PDCD4 transcriptional regulation target genes, clustered by fold change across sample type. Each row represents a gene, and each column represents a sample type. The color legend represents Bic vs TTX log2FC with red representing upregulation, green representing downregulation, and black indicating a log2FC of zero. **B)** GO analysis −log10 false discovery rate (FDR; circles) and percent of genes (bars) in terms from Biological Process (top four terms) and Cellular Compartment (bottom four terms) analyses (Ashburner et al., 2000; Carbon et al., 2019; Mi et al., 2019). Data from putative PDCD4 target genes (91 genes) are shown in red and for comparison, data from other CHX-insensitive activity-dependent genes (368 genes) are shown in blue. Select GO terms are shown for simplicity (see **Table S3** for top 15 GO terms by FDR for both groups of genes). **C)** Diagram depicting a generic synapse and synaptic proteins. The labeled synaptic proteins are encoded by putative PDCD4 target genes (gene name indicated in parenthesis alongside protein). The activity-dependent changes in expression of these genes are inhibited by degradation-resistant PDCD4. The presynaptic terminal is shown above with neurotransmitter-loaded synaptic vesicles, and the postsynaptic terminal is shown below with neurotransmitter receptors in the postsynaptic membrane (one receptor is shown anchored to an unlabeled gray PSD-95 protein). Arrow next to gene name illustrates the direction of activity-dependent differential expression and dashed line with bar illustrates the suppression of this activity-dependent change in the PDCD4 S71A samples.

## Discussion

In this study, we implemented a proximity-ligation assay to systematically characterize changes in the nuclear proteome triggered by neuronal stimulation. While advances in transcriptomic technologies have enabled the identification of genes that undergo activity-dependent changes in expression (Brigidi et al., 2019; Chen et al., 2017; Fernandez-Albert et al., 2019; Tyssowski et al., 2018), the systematic identification of proteins that undergo changes in subcellular localization and/or stability has been more challenging. Our results provide the first, to our knowledge, unbiased characterization of the population of proteins that undergo changes in nuclear abundance following neuronal silencing and/or glutamatergic stimulation, and does so in a manner that is independent of translation or transcription. We detected activity-dependent changes in the known nucleocytoplasmic shuttling proteins, CRTC1 and HDAC 4/5 (Ch’ng et al., 2012; Chawla et al., 2003), demonstrating the validity of this neuron-specific, subcellular compartment-specific assay.

The tumor suppressor protein PDCD4 was among the novel proteins we identified as undergoing activity-dependent changes in nuclear concentration. Low concentrations of PDCD4 have been reported to correlate with invasion, proliferation, and metastasis of many types of cancers (Allgayer, 2010; Chen et al., 2003; Wang & Yang, 2018; Wei et al., 2012). The activity-dependent downregulation of PDCD4 in neurons is reminiscent of the concept of “memory suppressor genes” (Abel & Kandel, 1998), genes that act as inhibitory constraints on activity-dependent neuronal plasticity. By analogy to its function during cancer metastases, decreases in PDCD4 in neurons would function to enable experience-dependent neuronal growth and remodeling. Dysregulated PDCD4 concentrations have also been reported to underlie a variety of metabolic disorders, including polycystic ovary syndrome, obesity, diabetes, and atherosclerosis, highlighting the critical role PDCD4 plays in regulating gene expression in multiple cell types (Lu et al., 2020). Despite being highly expressed, few studies have examined the function of PDCD4 in neurons (Di Paolo et al., 2020; Li et al., 2020; Narasimhan et al., 2013), and as far as we are aware, no previous study has identified a role for PDCD4 in activity-dependent gene regulation in neurons.

In investigating how neuronal activity decreased PDCD4 protein concentrations, we showed that it occurred via proteasome-mediated degradation. The finding that the nuclear export inhibitor LMB did not block the activity-dependent decrease of nuclear PDCD4 demonstrated that nuclear PDCD4 is degraded without leaving the nucleus. Many examples of activity-dependent degradation of proteins *within the cytoplasm* have been reported in neurons (Banerjee et al., 2009; Hegde et al., 1993; Jarome et al., 2011), but fewer cases of activity-dependent degradation of proteins *within the nucleus* have been described (Bayraktar et al., 2020; Kravchick et al., 2016; Upadhya et al., 2004). Nonetheless, the nucleus contains machinery for proteasome-mediated degradation and there are numerous examples of proteins that are degraded by the nuclear proteasome in non-neuronal cells, including transcriptional regulators and cell-cycle proteins (von Mikecz, 2006). Neuronal nuclei have also been shown to contain machinery for proteasome-mediated degradation (Mengual et al., 1996) and exhibit proteasomal activity, albeit with less activity than is present in the cytoplasm (Tydlacka et al., 2008; Upadhya et al., 2006).

Degradation of PDCD4 is regulated by phosphorylation of PDCD4 by the kinases S6K and PKC (Dorrello et al., 2006; Matsuhashi et al., 2014, 2019; Nakashima et al., 2010; Schmid et al., 2008). In our experiments, we found that a phospho-incompetent serine-to-alanine mutation at Ser71 prevented the activity-dependent degradation of PDCD4, and that phosphorylation by PKC, but not S6K was required for this activity-dependent degradation. This result is consistent with previous studies demonstrating that either S6K or PKC is required for PDCD4 phosphorylation depending on the signaling pathway, with EGF treatment requiring mTOR/S6K for PDCD4 degradation and TPA treatment requiring PKC (Matsuhashi et al., 2019). The stimulus-specific requirement of either S6K or PKC for PDCD4 degradation raises the interesting possibility that different types of neuronal stimulation could trigger PDCD4 degradation via distinct signaling pathways. Supporting this idea, two studies in neurons have suggested that PDCD4 degradation may be mediated by S6K (Di Paolo et al., 2020; Li et al., 2020), while our study demonstrated that PDCD4 degradation was mediated by PKC but not S6K. PKC is typically activated at the cell surface (Gould & Newton, 2008), however, it can translocate from the cytoplasm to the nucleus after activation and can be activated directly in the nucleus (Lim et al., 2015; Martelli et al., 2006), where it phosphorylates nuclear targets including histones and transcription factors (Lim et al., 2015; Martelli et al., 2006).

PDCD4 has been well-characterized as a translational repressor in cancer cells (Wang et al., 2017; Wedeken et al., 2011; Yang et al., 2004), and more recently in studies in neurons (Di Paolo et al., 2020; Li et al., 2020; Narasimhan et al., 2013). PDCD4 binds to the RNA helicase eIF4A and inhibits translation of mRNAs, particularly those with highly structured 5’ UTRs, including the cell cycle regulator *p53* (Wedeken et al., 2011), the cell growth regulator *Sin1* (Wang et al., 2017), and, as recently discovered in neurons, the neurotrophic growth factor *Bdnf* (Li et al., 2020). Further indicative of a role for PDCD4-mediated translational regulation in neurons, a recent study identified 267 putative translational targets of PDCD4 in PC12 cells, with decreases in PDCD4 leading to increased axonal growth in PC12 cells and in cultured primary cortical neurons (Di Paolo et al., 2020).

While the role of PDCD4 as a translational repressor has been well studied, the role of PDCD4 in the nucleus is less well-characterized, even though the protein is predominantly localized in the nucleus of many cells (Böhm et al., 2003). Our study, however, identified PDCD4 as a protein that underwent a decrease in nuclear concentration in response to Bic stimulation, and we detected greater activity-dependent decrease of PDCD4 in the nucleus than in the cytoplasm, suggesting a possible role for PDCD4 in the nucleus. Previous studies have indicated that, in non-neuronal cells, PDCD4 has been shown to inhibit AP-1-dependent transcription, although it is unclear whether this is a direct role in the nucleus (Bitomsky et al., 2004) or an indirect role regulating the translation of signaling proteins in the cytoplasm (Yang et al., 2006). PDCD4 has also been shown to bind to the transcription factors CSL (Jo et al., 2016) and TWIST1 (Shiota et al., 2009) and inhibit their transcriptional activity. In our study, we found 91 genes that are putative targets of PDCD4, including genes encoding proteins that are important for synaptic function. These findings suggest that the proteasome-mediated degradation of PDCD4 in the nucleus is important for regulating activity-dependent transcription following neuronal stimulation.

Taken together, our findings illustrate the utility of proximity-ligation assays in identifying activity dependent changes in the proteome of subcellular neuronal compartments and point to the array of cell biological mechanisms by which activity can regulate the neuronal proteome. They also focus attention on the tumor suppressor gene PDCD4 as a critical regulator of activity-dependent gene expression in neurons, highlighting a role for PDCD4 in regulating the transcription of genes involved in synapse formation, remodeling, and transmission. Future investigation of the mechanisms by which PDCD4 regulates transcription of these genes provides a means of characterizing the role of PDCD4 as a transcriptional regulator in addition to its previously well-characterized role as a translational inhibitor (Wang & Yang, 2018). Such studies also promise to deepen our understanding of the specific cell and molecular biological mechanisms by which experience alters gene expression in neurons to enable the formation and function of neural circuits.

## Methods

### RESOURCE AVAILABILITY

#### Lead Contact

Further information and requests for resources and reagents should be directed to and will be fulfilled by the Lead Contact, Kelsey Martin.

#### Materials Availability

Plasmids generated in this study are available upon request from the Lead Contact.

#### Data Availability

The published article includes the mass spectrometry data generated during this study. The full RNA sequencing data is available online (GEO Accession Number: GSE163127).

### EXPERIMENTAL MODEL DETAILS

#### Primary Neuronal Cultures

All experiments were performed using approaches approved by the UCLA Animal Research Committee. Forebrain from postnatal day 0 Sprague-Dawley rats (Charles River) was dissected in cold Hanks’ Balanced Salt Solution (HBSS, Thermo Fisher) supplemented with 10 mM HEPES buffer and 1 mM sodium pyruvate. Sex was not determined and tissue from male and female pups were pooled. The tissue was chopped finely and digested in 1x trypsin solution (Thermo Fisher) in HBSS (supplemented with 120 μg/mL DNase and 1.2 mM CaCl2) for 15 minutes at 37°C. The tissue was washed and triturated in Dulbecco’s Modified Eagle Medium (Thermo Fisher) + 10% Fetal Bovine Serum (Omega Scientific) before plating on poly-DL-lysine (PDLL)-coated (0.1 mg/mL, Sigma) 10 cm dishes or 24-well plates containing acid-etched PDLL-coated coverslips (Carolina Biologicals). Neurons were plated at a density of 1 forebrain per 10 cm dish (for mass spectrometry experiments) or 1/2 forebrain per entire 24-well plate (for immunocytochemistry, RNA-seq, and western blot experiments). Neurons were cultured in Neurobasal-A (Thermo Fisher) supplemented with 1x B-27 (Thermo Fisher), 0.5 mM glutaMAX (Thermo Fisher), 25 μM monosodium glutamate (Sigma), and 25 μM β-mercaptoethanol (Sigma) and incubated at 37°C, 5% CO2. When applicable, neurons were transfected with plasmids using Lipofectamine 2000 (Thermo Fisher) according to manufacturer’s instructions at days in vitro (DIV) 2, or transduced with AAV at DIV 13. All experiments were performed at DIV 20.

### METHOD DETAILS

#### Generation of Plasmids and AAV

To create hSyn NLS-APEX2-EGFP-NLS, APEX2 was amplified from the pcDNA3 APEX2-NES plasmid (gift from Alice Ting, Addgene plasmid #49386) with three sequential sets of primers to add SV40 NLS to both the N-terminus and C-terminus of APEX2 (primer sets 1-3, **Table S4**). The design of using NLS on both sides of APEX2 was based on the design of Cas9-NLS (Swiech et al., 2015). NLS-APEX2-NLS was then inserted into the pAAV-hSyn-EGFP plasmid (gift from Bryan Roth, Addgene plasmid #50465) between the BamHI and EcoRI sites, replacing the EGFP insert. The final hSyn NLS-APEX2-EGFP-NLS construct was created by amplifying hSyn NLS-APEX2-NLS (primer set 4, **Table S4**) and EGFP (primer set 5, **Table S4**), and joining the two products at NheI and SacI.

To create hSyn PDCD4-HA, rat PDCD4 was amplified from cultured neuron cDNA (primer set 6, **Table S4**) with a C-terminal HA tag, and then inserted into the pAAV-hSyn-EGFP plasmid between NcoI and EcoRI, replacing the EGFP insert. The S71A mutation was created using site-directed mutagenesis (services by Genewiz) to mutate serine 71 (TCT) to alanine (GCT). The hSyn V5-PDCD4-HA construct was created by adding a V5 tag to the N-terminus of PDCD4-HA using PCR-based mutagenesis (services by Genewiz).

AAV9 was generated for APEX2-NLS, PDCD4-HA WT, and PDCD4-HA S71A at Penn Vector Core.

#### Pharmacological Treatments

Neurons were pre-incubated with cycloheximide (CHX, 60 μM, Sigma) for 15 min, leptomycin B (LMB, 10 nM, Sigma) for 30 min, LY2584702 (1 μM, Cayman Chemical) or Ro-31-8425 (5 μM, Sigma) for 1 hr, or epoxomicin (5 μM, Enzo Life Sciences), bortezomib (10 μM, APExBIO), or MLN4924 (50 nM, APExBIO) for 2 hrs before the start of each respective experiment and remained in the media throughout the duration of each experiment, incubated at 37 °C. For neurons treated with inhibitors dissolved in DMSO (epoxomicin, bortezomib, MLN4924, LY2584702, and Ro-31-8425), the final DMSO concentration in the media was 0.1% or less. For neurons treated with LMB, the final methanol concentration in the media was 0.08%. All control groups received an equivalent concentration of vehicle (DMSO or methanol). To silence the neurons, 1 μM tetrodotoxin (TTX, Tocris) was applied to the neurons for 1 hr. To stimulate the neurons, 40 μM (-)-bicuculline methiodide (Bic, Tocris) was applied to the neurons for 1 hr unless otherwise stated.

For KCl stimulations, neurons were pre-incubated with standard Tyrode’s solution (140 mM NaCl, 10 mM HEPES, 5 mM KCl, 3 mM CaCl_2_, 1 mM MgCl_2_, 20 mM glucose, pH 7.4) containing 1 μM TTX for 15 min at room temperature, and then stimulated for 5 min with 40 mM KCl isotonic Tyrode’s solution containing TTX. Control cells remained in the standard Tyrode’s solution containing TTX throughout the experiment.

Control data from LMB (**Fig 3A, Fig S2E**) and CHX (**Fig S2A-B**) experiments were combined to generate the data shown in **Fig 2B-C**. Two of the bortezomib experiments were performed concurrently with two of the MLN4924 experiments, and so these experiments partially share control data in **Fig 3C-D** and **Fig S2G-H**. One of the LY2584702 experiments was performed concurrently with one of the bortezomib experiments, and so **Fig 4E** and **Fig S3B** partially shares control data with **Fig 3C** and **Fig S2G**.

#### APEX2 Proximity Biotinylation, Streptavidin Pulldown, and On-Bead Tryptic Digestion

For APEX2 mass spectrometry experiments, 3 biological replicates (sets of cultures) were prepared, with 3 samples in each replicate (APEX + Bic, APEX + TTX, and No APEX). Neurons were TTX-silenced or Bic-stimulated for 1 hour in the presence of CHX (as above). During the final 30 minutes of the treatment, neurons were incubated with 500 μM biotin-phenol (APExBIO) at 37 °C. During the final 1 minute, labeling was performed by adding H_2_O_2_ to a final concentration of 1 mM. To stop the labeling reaction, neurons were washed three times in large volumes of quencher solution (phosphate-buffered saline with 10 mM sodium azide, 10 mM sodium ascorbate, and 5 mM Trolox).

Neurons were lysed with RIPA (50 mM Tris, 150 mM NaCl, 0.1% SDS, 0.5% sodium deoxycholate, 1% Triton X-100, pH 7.5) containing protease inhibitor cocktail (Sigma), phosphatase inhibitor cocktail (Sigma), and quenchers. Lysates were treated with benzonase (200 U/mg protein, Sigma) for 5 min and then clarified by centrifugation at 15,000 g for 10 min. Samples were concentrated with Amicon centrifugal filter tubes (3K NMWL, Millipore) to at least 1.5 mg/mL protein and quantified using Pierce 660 nm protein assay kit Thermo Fisher).

For each sample, 2 mg of lysate was incubated with 220 μL Pierce streptavidin magnetic beads (Thermo Fisher) for 60 min at room temperature. Samples were washed twice with RIPA, once with 1M KCl, once with 0.1 M sodium carbonate, once with 2 M Urea 10 mM Tris-HCl pH 8.0, and twice with RIPA according to (Hung et al., 2016).

The streptavidin beads bound by biotinylated proteins were then washed three times with 8 M Urea 100 mM Tris-HCl pH 8.5 and three times with pure water, and then the samples were resuspended in 100 μl 50 mM TEAB. Samples were reduced and alkylated by sequentially incubating with 5 mM TCEP and 10 mM iodoacetamide for 30 minutes at room temperature in the dark on a shaker set to 1000 rpm. The samples were incubated overnight with 0.4 μg Lys-C and 0.8 μg trypsin protease at 37° C on a shaker set to 1000 rpm. Streptavidin beads were removed from peptide digests, and peptide digests were desalted using Pierce C18 tips (100 μl bead volume), dried, and then reconstituted in water.

#### Tandem Mass Tag (TMT) Labeling

The desalted peptide digests were labeled by TMT reagents according to the manufacturer’s instructions (TMT10plex™ Isobaric Label Reagent Set, catalog number). Essentially, peptides were incubated with acetonitrile reconstituted TMT labeling reagent for 1 hour and then quenched by adding hydroxylamine. Sample-label matches are: NoAPEX replicate #1 labeled with TMT126, APEX+Bic replicate #1 labeled with TMT127N, APEX+TTX replicate #1 labeled with TMT127C, APEX+TTX replicate #2 labeled with TMT128N, NoAPEX replicate #2 labeled with TMT128C, NoAPEX replicate #3 labeled with TMT129N, APEX+Bic replicate #2 labeled with TMT129C, APEX+Bic replicate #3 labeled with TMT130N, APEX+TTX replicate #3 labeled with TMT130C. Labeled samples were then combined, dried and reconstituted in 0.1% TFA for high pH reversed phase fractionation.

#### High pH Reversed Phase Fractionation

High pH reversed phase fractionation was performed according to the manufacturer’s instructions (Pierce High pH Reversed-Phase Peptide Fractionation Kit). Essentially, peptides were bound to the resin in the spin column and then eluted by stepwise incubations with 300 μl of increasing acetonitrile concentrations. The eight fractions were combined into four fractions (fractions 1 & 5, 2 & 6, 3 & 7, 4 & 8). Fractions were then dried by vacuum centrifugation and reconstituted in 5% formic acid for mass spectrometry analysis.

#### LC-MS Data Acquisition

A 75 μm x 25 cm custom-made C18 column was connected to a nano-flow Dionex Ultimate 3000 UHPLC system. A 140-minute gradient of increasing acetonitrile (ACN) was delivered at a 200 nL/min flow rate as follows: 1 – 5.5% ACN phase from minutes 0 – 5, 5.5 – 27.5% CAN from minutes 5 – 128, 27.5 – 35% ACN from minutes 128 – 135, 35 – 80% ACN from minutes 135 – 136, 80% ACN hold from minutes 136 – 138 and then down to 1% ACN from minutes 138 – 140. An Orbitrap Fusion Lumos Tribrid mass spectrometer TMT-MS3-SPS method was used for data acquisition. Full MS scans were acquired at 120K resolution in Orbitrap with the AGC target set to 2e5 and a maximum injection time set to 50 ms. MS2 scans were collected in Ion Trap with Turbo scan rate after isolating precursors with an isolation window of 0.7 m/z and CID fragmentation using 35% collision energy. MS3 scans were acquired in Obitrap at 50K resolution and 10 synchronized selected precursor ions were pooled for each scan using 65% HCD energy for fragmentation. For data dependent acquisition, a 3-second cycle time was used to acquire MS/MS spectra corresponding to peptide targets from the preceding full MS scan. Dynamic exclusion was set to 30 seconds.

#### MS/MS Database Search

MS/MS database searching was performed using MaxQuant (1.6.10.43) (Cox & Mann, 2008) against the rat reference proteome from EMBL (UP000002494-10116 RAT, Rattus norvegicus, 21649 entries). The search included carbamidomethylation on as a fixed modification and methionine oxidation and N-terminal acetylation as variable modifications. The digestion mode was set to trypsin and allowed a maximum of 2 missed cleavages. The precursor mass tolerances were set to 20 and 4.5 ppm for the first and second searches, respectively, while a 20-ppm mass tolerance was used for fragment ions. Datasets were filtered at 1% FDR at both the PSM and protein-level. Quantification type was set to reporter ion MS3 with 10plex TMT option.

#### Statistical Inference in Mass Spectrometry Data

MSStatsTMT (1.4.1) (Huang et al., 2020) was used to analyze the MaxQuant TMT-MS3 data in the APEX2 proximity labeling experiment to statistically assess protein enrichment. TTX channels were used for MS run level normalization. The “msstats” method was then used for protein summarization. P-values for t-tests were corrected for multiple hypothesis testing using the Benjamini-Hochberg adjustment. We identified proteins that were enriched above the No APEX negative control using a log2FC > 3 and adjusted p-value < 0.05 cutoff above the No APEX condition. Of this protein list, we then identified proteins that were differentially expressed when comparing between Bic and TTX conditions using log2FC > 0.5 (for Bic) or log2FC < −0.5 (for TTX) with p-value < 0.05. It is important to note that we used a non-adjusted p-value cutoff when identifying candidate proteins that were differentially expressed between the TTX-silenced and Bic-stimulated conditions, because only HDAC4 and six other proteins had a significant adjusted p-value when using these cutoffs. Even CRTC1, a protein that has been shown to undergo activity-dependent changes in multiple studies (Ch’ng et al., 2012, 2015; Nonaka et al., 2014) and confirmed again in the present study, did not reach adjusted p-value significance, suggesting that we do not have the statistical power to detect certain activity-dependent changes. Because we used non-adjusted p-values to identify candidate proteins, it is especially important to experimentally validate any potential candidate protein.

#### Protein Extraction and Western Blot

Neurons were washed in Tyrode’s solution (140 mM NaCl, 10 mM HEPES, 5 mM KCl, 3 mM CaCl_2_, 1 mM MgCl_2_, 20 mM glucose, pH 7.4) and lysed with RIPA (50 mM Tris, 150 mM NaCl, 0.1% SDS, 0.5% sodium deoxycholate, 1% Triton X-100, pH 7.5) containing protease and phosphatase inhibitor cocktails (Sigma). Samples were clarified by centrifugation at 10,000 g for 10 min. Protein concentration was determined using the Pierce BCA protein assay kit (Thermo Fisher).

Protein lysates were boiled in loading buffer (10% glycerol, 1% SDS, 60 mM Tris HCl pH 7.0, 0.1 M DTT, 0.02% bromophenol blue) for 10 min at 95°C and run on at 8% polyacrylamide gel for 90 min at 120 V. Samples were wet-transferred onto a 0.2 μm nitrocellulose membrane for 16 hours at 40 mA. The membrane blocked with Odyssey Blocking Buffer (LI-COR) and incubated with primary antibodies: mouse HA (BioLegend #901513, 1:1000), mouse TUJ1 (BioLegend #801201, 1:1000), mouse S6 (CST #2317, 1:1000), rabbit phospho-S6 ser 235/236 (CST #4858, 1:2000) for 3-4 hours at room temperature or overnight at 4°C. The membrane was washed with TBST and incubated with secondary antibodies: anti-rabbit IRDye 800CW (1:10,000), anti-mouse IRDye 800CW (1:10,000), anti-mouse IRDye 680CW (1:10,000), IRDye 800CW Streptavidin (1:1,000) for 1 hour at room temperature. The membrane was imaged using the Odyssey Infrared imaging system (LI-COR). Western blots were quantified using the Image Studio (LI-COR) rectangle tool. The relative intensity of each band was calculated by normalizing to a loading control (TUJ1). Within each experiment, all values were normalized to the control (basal) sample.

#### Immunocytochemistry (ICC)

Neurons were fixed with 4% paraformaldehyde in phosphate-buffer saline (PBS) for 10 min, permeabilized in 0.1% triton X-100 in PBS for 5 min, and blocked in 10% goat serum in PBS for 1 hour. Neurons were incubated with primary antibodies: chicken MAP2 (PhosphoSolutions #1100-MAP2, 1:1000), rabbit PDCD4 (CST #9535, 1:600), mouse HA (BioLegend #901513, 1:1000), mouse V5 (Thermo Fisher #R960-25, 1:250), rabbit CRTC1 (Bethyl #A300-769, 1:1000), rabbit HDAC4 (CST #7628, 1:100), rabbit FOS (CST #2250, 1:500) for 3-4 hours at room temperature or overnight at 4°C. Neurons were washed with PBS, and incubated at 1:1000 with secondary antibodies: anti-chicken Alexa Fluor 647, anti-rabbit Alexa Fluor 555, anti-mouse Alexa Fluor 555, Streptavidin Alexa Fluor 555, and Hoechst 33342 stain for 1 hour at room temperature. Neurons were washed with PBS, and mounted on slides with Aqua-Poly/Mount (Polysciences) for confocal imaging.

#### Confocal Imaging

Samples were imaged using a Zeiss LSM 700 confocal microscope with a 40x oil objective and 405 nm, 488 nm, 555 nm, and 639 nm lasers. Identical image acquisition settings were used for all images within an experiment. For each image acquisition, the experimenter viewed the MAP2 and Hoechst channels to select a field-of-view, and was blind to the experimental channel (HA, PDCD4, etc.). For each coverslip, images were taken at multiple regions throughout the coverslip, and 2-3 coverslips were imaged per condition. Images were collected from at least 3 experimental replicates (sets of cultures), unless otherwise stated.

#### Image Analysis

ICC images were processed using ImageJ (Schindelin et al., 2012). An ImageJ macro was used to create regions of interest (ROIs) for neuronal nuclei. In brief, the Hoechst signal was used to outline the nucleus, and the MAP2 signal was used to select neurons and exclude non-neuronal cells. To create ROIs for neuronal cytoplasm, the cell body of each neuron was manually outlined using the MAP2 signal and then the nuclear ROI was subtracted from the total cell body ROI. The ROIs were used to calculate the mean intensity in the channel of interest (HA, PDCD4, etc.) for the nucleus and cytoplasm of each neuron. Within each ICC experimental replicate, the measured values from all ROIs were normalized to the median value of the control condition (basal). For experiments using transfected cells (V5 experiments), the measured intensity of each ROI was normalized to the co-transfection marker (nuclear GFP intensity), in order to normalize for differences in transfection efficiency between cells.

#### RNA Extraction, Library Preparation, RNA Sequencing, and Data Analysis

Samples were prepared from 3 biological replicates (sets of cultures), with 8 samples in each replicate (WT Bic, WT TTX, S71A Bic, S71A TTX, CHX WT Bic, CHX WT TTX, CHX S71A Bic, CHX S71A TTX). RNA was extracted from neuronal cultures using the RNeasy Micro Kit (Qiagen) according to manufacturer’s instructions. Libraries for RNA-Seq were prepared with Nugen Universal plus mRNA-Seq Kit (Nugen) to generate strand-specific RNA-seq libraries. Samples were multiplexed, and sequencing was performed on Illumina HiSeq 3000 to a depth of 25 million reads/sample with single-end 65 bp reads. Demultiplexing was performed using Illumina Bcl2fastq v2.19.1.403 software. The RNA-seq data discussed in this publication have been deposited in NCBI’s Gene Expression Omnibus (Edgar et al., 2002) and are accessible through GEO Series accession number GSE163127 (https://www.ncbi.nlm.nih.gov/geo/query/acc.cgi?acc=GSE163127). Reads were aligned to Rattus_norvegicus reference genome version Rnor_6.0 (rn6), and reads per gene was quantified by STAR 2.27a (Dobin et al., 2013) using Rnor_6.0 gtf file. We used DESeq2 (Love et al., 2014) to obtain normalized read counts and perform differential expression analysis, including batch correction for replicate number (**Table S2**). Putative PDCD4 target genes were identified by first focusing on genes which showed activity-dependent differential expression in both the presence and absence of CHX (CHX-insensitive activity-dependent genes; 459 genes after excluding 3 genes which showed differential expression in different directions +/− CHX). We then calculated the PDCD4 activity-dependent change by taking the difference between the activity-dependent fold change of PDCD4 WT and PDCD4 S71A samples and normalizing:

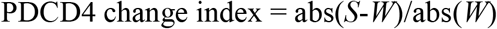

Where *S* is the PDCD4 S71A no CHX Bic vs TTX log2FC, and *W* is the PDCD4 WT no CHX Bic vs TTX log2FC. We defined putative PDCD4 target genes as those with a PDCD4 change index > 0.75. Motif analysis was performed using the `findMotifsGenome` command in HOMER (S. Heinz et al., 2010), using sequences from the TSS and upstream 500 bp as the promoter sequences for each gene. GO analysis was performed using the Gene Ontology Resource (Ashburner et al., 2000; Carbon et al., 2019) and PANTHER enrichment tools (Mi et al., 2019). Cartoon of putative PDCD4 targets was generated using BioRender.com.

#### RT-qPCR

As above, RNA was extracted from neuronal cultures using the RNeasy Micro Kit (Qiagen). cDNA was synthesized from 500 ng RNA using SuperScript III First-Strand Synthesis System (Thermo Fisher) with random hexamers. A “No Reverse Transcriptase” sample was also prepared as a negative control. RT-qPCR was performed on the CFX Connect Real-Time System (Bio-Rad) using PowerUp SYBR Green Master Mix (Applied BioSystems). Primer pairs were designed for two housekeeping genes (*Hprt*, *Gapdh*) and two candidate genes (*Scd1*, *Thrsp*) using Primer3Plus (Untergasser et al., 2012) and NCBI Primer-BLAST (Ye et al., 2012) (**Table S4**). RT-qPCR was performed on 6 sets of cultures, with technical triplicate reactions for each sample. For each gene, relative quantity was calculated using the formula: E^(ΔCt), where E was calculated from the primer efficiencies (E≈2) and ΔCt was calculated using Ct TTX– Ct Bic. Relative gene expression was calculated by normalizing the relative quantity of the gene of interest to the relative quantity of the housekeeping genes *Hprt* and *Gapdh*: (E _gene_)^ΔCt gene^ / GeoMean[(E _HPRT_)^ΔCt HPRT^, (E _GAPDH_)^ΔCt GAPDH^].

### QUANTIFICATION AND STATISITCAL ANALYSIS

For ICC experiments, the quantification of signal intensity is displayed in violin plots using GraphPad Prism. The medians are indicated with thick lines and the quartiles are indicated with thin lines. “n” refers to the number of neurons in each condition, and all individual data points were plotted on the graphs. Our sample sizes were not pre-determined. A non-parametric statistical test (Mann-Whitney U test) was used to calculate statistical significance because our data were not normally distributed, as indicated by the violin plots. A Bonferroni correction was used to adjust for multiple hypothesis testing. Statistical significance is indicated by *p < 0.05, **p < 0.01, ***p < 0.001, and ****p < 0.0001. For Main Figures, the sample sizes are indicated in the figure legends and the medians, statistical tests, and p-values are indicated in the results section. For Supplementary Figures, the sample sizes, medians, statistical tests, and p-values are all indicated in the figure legends.

For RT-qPCR experiments, all data points were displayed using GraphPad Prism, with solid lines indicating the median values. “n” refers to the biological replicates (sets of cultures), and all data points were plotted on the graphs. The Mann-Whitney U test (Prism) was used to calculate statistical significance. Statistical significance is indicated by *p < 0.05 and **p < 0.01. The sample size, medians, statistics tests, and p-values are all indicated in the supplementary figure legend.

For western blot experiments, all data points were displayed using GraphPad Prism, with solid lines indicating the median values. “n” refers to the biological replicates (sets of cultures), and all data points were plotted on the graphs. The medians are indicated in the results section and the sample sizes are indicated in the figure legends.

## Acknowledgements

We thank Sylvia Neumann, Marika Watanabe, and Emilie Marcus for their comments on the manuscript, and members of the Martin lab for helpful discussion. RNA sequencing was performed at the Technology Center for Genomics & Bioinformatics (TCGB) at UCLA. This work was supported by NIH grants R01MH077022 to K.C.M. and NRSA F31MH113310 to W.A.H.

## Author Contributions

Conceptualization: W.A.H. and K.C.M.

Methodology: W.A.H., W.D., J.A.W., and K.C.M

Investigation: W.A.H. (APEX2 and PDCD4 neuron experiments), W.D. (mass spectrometry), and J.M.A. (RNA-seq data analysis)

Writing – Original Draft: W.A.H., J.M.A, and K.C.M.

Writing – Review & Editing: W.A.H., J.M.A., and K.C.M.

Supervision: J.A.W., J.M.A., and K.C.M.

## Supplemental Information

**Figure S1.**
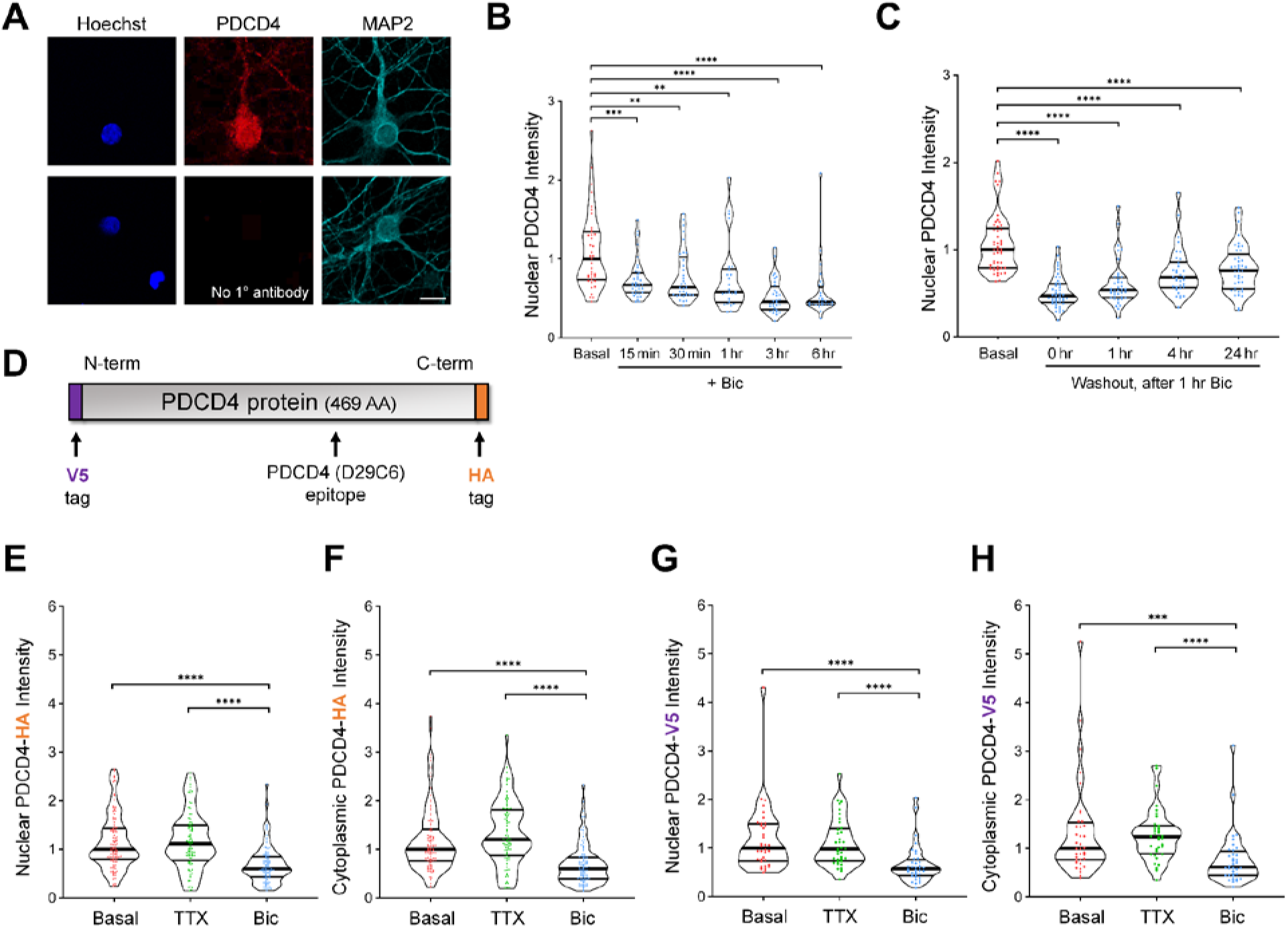
Relating to Figure 2: Neuronal stimulation decreases PDCD4 protein concentration in the nucleus and cytoplasm of neurons. **A)** Top: immunocytochemistry (ICC) of endogenous PDCD4 protein. Bottom: negative control (no primary antibody). Scale bar = 10 μm. **B)** Violin plots of normalized nuclear PDCD4 ICC intensity after varying durations of Bic stimulation. Basal n = 39, Bic 15 min n = 36, 30 min n = 33, 1 hr n = 26, 3 hr n = 32, 6 hr n = 28 cells, from 1 set of cultures. Basal median = 1.00, Bic 15 min median = 0.67, 30 min median = 0.64, 1 hr median = 0.58, 3 hr median = 0.46, 6 hr median = 0.46. Basal vs Bic 15 min p = 0.0005, Basal vs Bic 30 min p = 0.007, Basal vs Bic 1 hr p = 0.0035, Basal vs Bic 3 hr p < 0.0001, Basal vs Bic 6 hr p < 0.0001. **C)** Violin plots of normalized nuclear PDCD4 ICC intensity at various timepoints after removal of a 1 hr Bic stimulation. Basal n = 49, Washout 0 hr n = 44, 1 hr n = 40, 4 hr n = 34, 24 hr n = 41 cells, from 1 set of cultures. Basal median = 1.00, Washout 0 hr median = 0.47, 1 hr median = 0.54, 4 hr median = 0.68, 24 hr median = 0.76. Basal vs 0 hr p < 0.0001, Basal vs 1 hr p < 0.0001, Basal vs 4 hr p < 0.0001, Basal vs 24 hr < 0.0001. **D)** PDCD4 protein, with locations of V5 tag, HA tag, and PDCD4 (D29C6) epitope (recognized by the PDCD4 Cell Signaling Technology antibody used in this study). **E)** Violin plots of normalized nuclear HA ICC intensity in neurons transduced with PDCD4-HA AAV. Basal n = 107, TTX n = 88, Bic n = 102 cells, from 3 sets of cultures. Basal median = 1.00, TTX median = 1.116, Bic median = 0.5972. Basal vs Bic p < 0.0001, TTX vs Bic p < 0.0001. **F)** Violin plots of normalized cytoplasmic HA ICC intensity in the same cells as in C. Basal median = 1.00, TTX median = 1.203, Bic median = 0.5983. Basal vs Bic p < 0.0001, TTX vs Bic p < 0.0001. **G)** Violin plots of normalized nuclear V5 ICC intensity in neurons transfected with V5-PDCD4 plasmid and co-transfected with GFP plasmid as a transfection marker. Basal n = 36, TTX n = 36, Bic n = 36 cells, from 2 sets of cultures. For each cell, the nuclear V5 intensity was normalized to the nuclear GFP intensity, in order to normalize for differences in transfection efficiency between cells. Basal median = 1.00, TTX median = 0.9810, Bic median = 0.5760. Basal vs Bic p < 0.0001, TTX vs Bic p < 0.0001. **H)** Violin plots of normalized cytoplasmic V5 ICC intensity in the same cells as in E. For each cell, the cytoplasmic V5 intensity was normalized to the nuclear GFP intensity, in order to normalize for differences in transfection efficiency between cells. Basal median = 1.00, TTX median = 1.237, Bic median = 0.6167. Basal vs Bic p = 0.0002, TTX vs Bic p < 0.0001. Statistical significance is indicated by ***p < 0.001 and ****p < 0.0001, from Mann-Whitney U test with Bonferroni correction.

**Figure S2.**
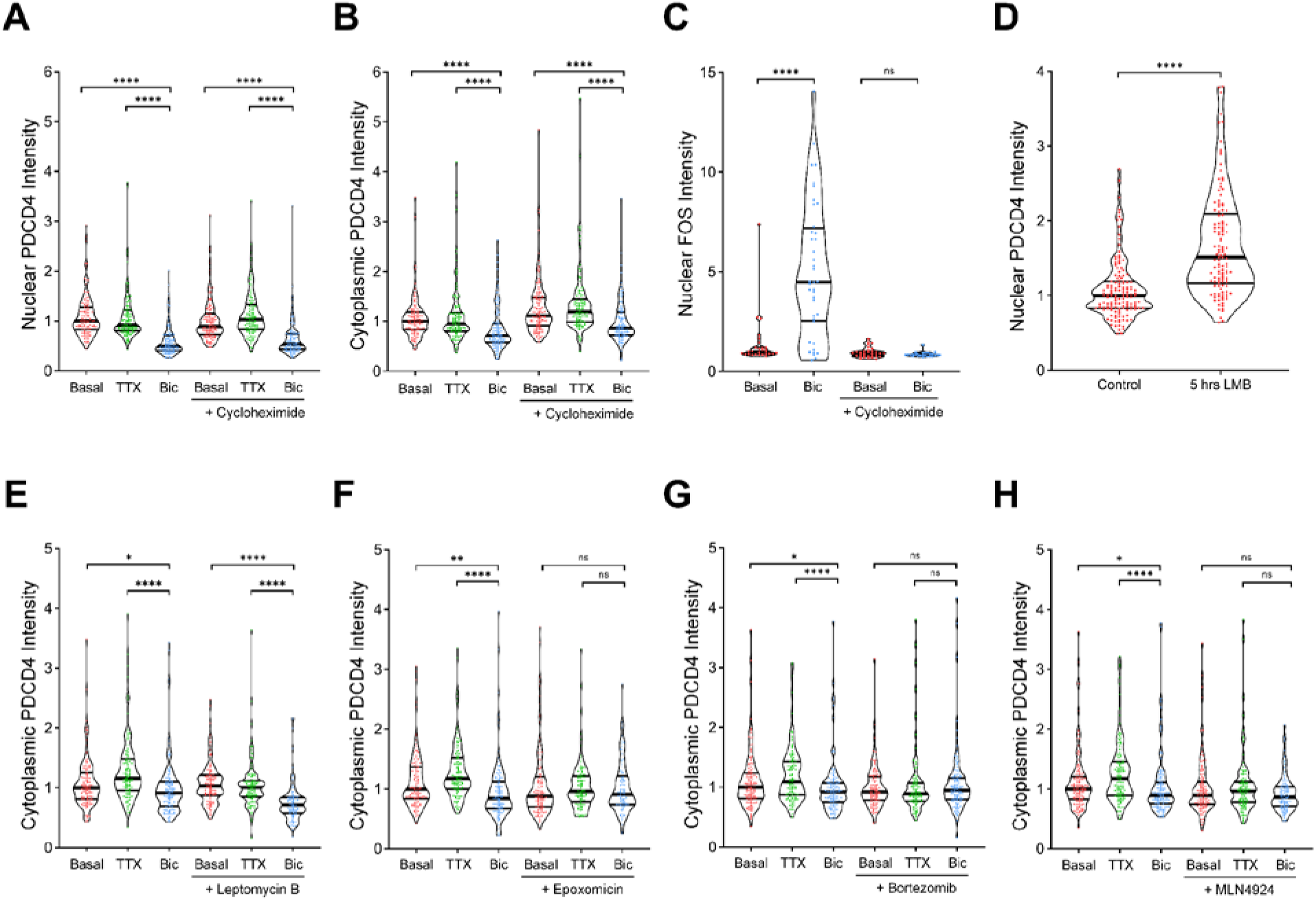
Relating to Figure 3: PDCD4 undergoes proteasome-mediated degradation –not nuclear export– following neuronal stimulation. **A)** Violin plots of normalized nuclear PDCD4 immunocytochemistry (ICC) intensity. Basal n = 117, TTX n = 118, Bic n = 109, CHX-Basal n = 123, CHX-TTX n = 120, CHX-Bic n = 104 cells, from 3 sets of cultures. Basal median = 1.00, TTX median = 0.9119, Bic median = 0.4924, CHX-Basal median = 0.8890, CHX-TTX median = 1.033, CHX-Bic median = 0.5375. Basal vs Bic p < 0.0001, TTX vs Bic p < 0.0001, CHX-Basal vs CHX-Bic p < 0.0001, CHX-TTX vs CHX-Bic p < 0.0001. **B)** Violin plots of normalized cytoplasmic PDCD4 ICC intensity in the same cells as in A. Basal median = 1.00, TTX median = 0.9501, Bic median = 0.7138, CHX-Basal median = 1.112, CHX-TTX median = 1.191, CHX-Bic median = 0.8626. Basal vs Bic p < 0.0001, TTX vs Bic p < 0.0001, CHX-Basal vs CHX-Bic p < 0.0001, CHX-TTX vs CHX-Bic p < 0.0001. **C)** Violin plots of normalized nuclear FOS ICC intensity. Basal n = 28, Bic n = 40 cells, CHX-Basal n = 32, CHX-Bic n = 26 cells, from 1 set of cultures. Basal median = 1.00, Bic median = 4.484, CHX-Basal median = 0.8941, CHX-Bic median = 0.8307. Basal vs Bic p < 0.0001, CHX-Basal vs CHX-Bic p = 0.3064. **D)** Violin plots of normalized nuclear PDCD4 ICC intensity. Control n = 137, LMB n = 122 cells, from 3 sets of cultures. Control median = 1.00, LMB median = 1.513. Control vs LMB p < 0.0001. **E)** Violin plots of normalized cytoplasmic PDCD4 ICC intensity in the same cells as in **Fig 3A**. Basal median = 1.00, TTX median = 1.158, Bic median = 0.9127, LMB-Basal median = 1.031, LMB-TTX median = 1.005, LMB-Bic median = 0.7122. Basal vs Bic p = 0.034, TTX vs Bic p < 0.0001, LMB-Basal vs LMB-Bic p < 0.0001, LMB-TTX vs LMB-Bic p < 0.0001. **F)** Violin plots of normalized cytoplasmic PDCD4 ICC intensity in the same cells as in **Fig 3B**. Basal median = 1.00, TTX median = 1.174, Bic median = 0.8439, Epox-Basal median = 0.8789, Epox-TTX median = 0.9596, Epox-Bic median = 0.9077. Basal vs Bic p = 0.001, TTX vs Bic p < 0.0001, Epox-Basal vs Epox-Bic p = 1, Epox-TTX vs Epox-Bic p = 0.8258. **G)** Violin plots of normalized cytoplasmic PDCD4 ICC intensity in the same cells as in **Fig 3C**. Basal median = 1.00, TTX median = 1.093, Bic median = 0.9226, Bort-Basal median = 0.9229, Bort-TTX median = 0.8904, Bort-Bic median = 0.9472. Basal vs Bic p = 0.0156, TTX vs Bic p < 0.0001, Bort-Basal vs Bort-Bic p = 1, Bort-TTX vs Bort-Bic p = 0.3544. **H)** Violin plots of normalized cytoplasmic PDCD4 ICC intensity in the same cells as in **Fig 3D**. Basal median = 1.00, TTX median = 1.173, Bic median = 0.8955, MLN-Basal median = 0.8940, MLN-TTX median = 0.9603, MLN-Bic median = 0.8633. Basal vs Bic p = 0.0324, TTX vs Bic p < 0.0001, MLN-Basal vs MLN-Bic p = 0.6294, MLN-TTX vs MLN-Bic p = 0.11. Statistical significance is indicated by *p < 0.05, **p < 0.01, and ****p < 0.0001, from Mann-Whitney U test with Bonferroni correction.

**Figure S3.**
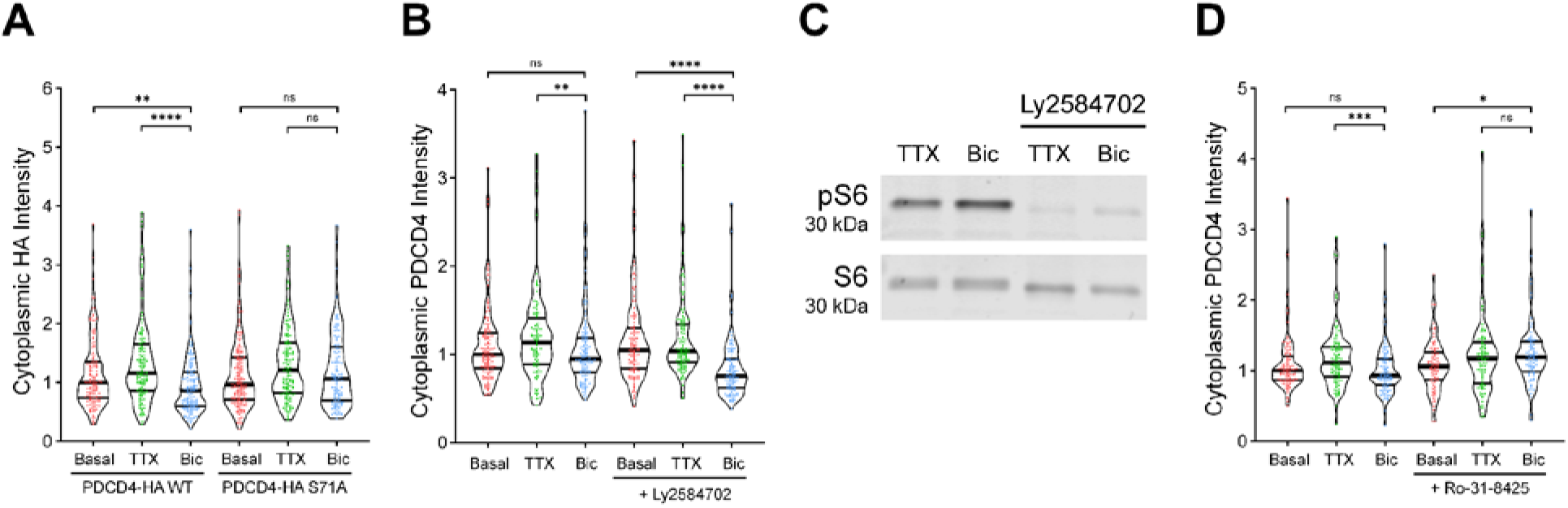
Relating to Figure 4: PDCD4 S71A mutation and PKC inhibition prevent the activity-dependent decrease of PDCD4. **A)** Violin plots of normalized cytoplasmic HA immunocytochemistry (ICC) intensity in the same cells as in **Fig 4C**. WT-Basal median = 1.00, WT-TTX median = 1.161, WT-Bic median = 0.8621, S71A-Basal median = 0.9676, S71A-TTX median = 1.209, S71A-Bic median = 1.063. WT-Basal vs WT-Bic p = 0.0012, WT-TTX vs WT-Bic p < 0.0001, S71A-Basal vs S71A-Bic p = 0.6704, S71A-TTX vs S71A-Bic p = 0.2012. **B)** Violin plots of normalized cytoplasmic PDCD4 ICC intensity in the same cells as in **Fig 4E**. Basal median = 1.00, TTX median = 1.14, Bic median = 0.95, LY-Basal median = 1.05, LY-TTX median = 1.04, LY-Bic median = 0.76. Basal vs Bic p = 0.4742, TTX vs Bic p = 0.0088, LY-Basal vs LY-Bic p < 0.0001, LY-TTX vs LY-Bic p < 0.0001. **C)** Western blot of protein lysates from neurons treated with or without LY2584702, from 1 set of cultures. Western blot was stained with antibodies for phospho-S6 (ser 235/236) and total S6 to confirm that LY2584702 inhibits S6K activity. **D)** Violin plots of normalized cytoplasmic PDCD4 ICC intensity in the same cells as in **Fig 4F**. Basal median = 1.00, TTX median = 1.114, Bic median = 0.9326, Ro-Basal median = 1.059, Ro-TTX median = 1.172, Ro-Bic median = 1.195. Basal vs Bic p = 0.2592, TTX vs Bic p = 0.0008, Ro-Basal vs Ro-Bic p = 0.0118, Ro-TTX vs Ro-Bic p = 0.9892. Statistical significance is indicated by *p < 0.05, **p < 0.01, ***p < 0.001, and ****p < 0.0001, from Mann-Whitney U test with Bonferroni correction.

**Figure S4.**
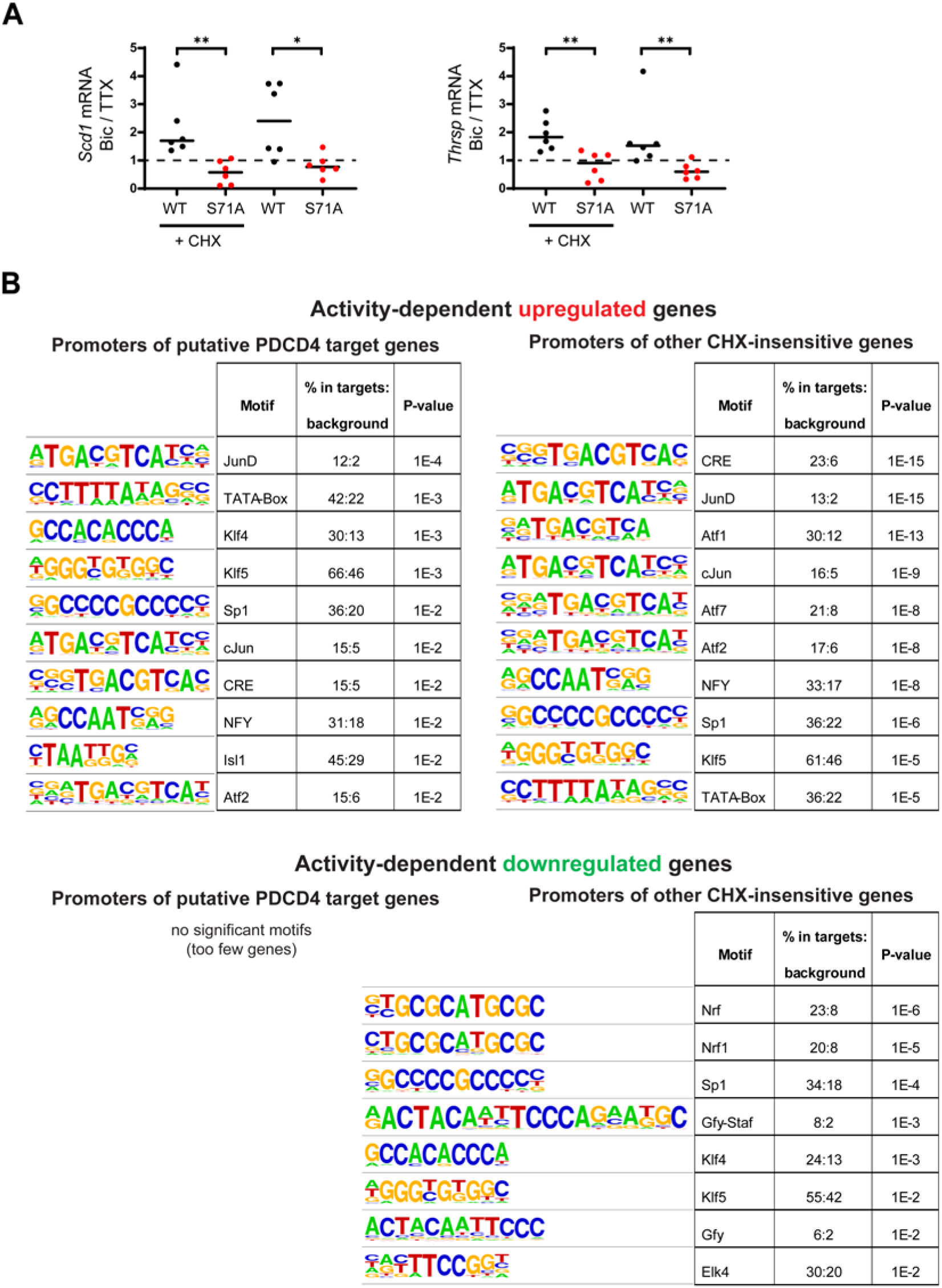
Relating to Figures 5–6: Stimulus-induced degradation of PDCD4 regulates the expression of neuronal activity-dependent genes. **A)** RT-qPCR of putative PDCD4 target genes, *Scd1* and *Thrsp*, from TTX-silenced and Bic-stimulated neurons that were transduced with PDCD4 WT or S71A, from 6 sets of cultures. Samples were normalized using two housekeeping genes, *Hprt* and *Gapdh*. The abundance of the target gene in each Bic sample was normalized to its respective TTX sample. *Scd1* WT CHX median = 1.702, *Scd1* S71A CHX median = 0.5776, *Scd1* WT median = 2.401, *Scd1* S71A median = 0.7672, *Thrsp* WT CHX median = 1.826, *Thrsp* S71A CHX median = 0.9080, *Thrsp* WT median = 1.522, *Thrsp* S71A median = 0.5995. *Scd1* CHX WT vs CHX S71A p = 0.0022, *Scd1* WT vs S71A p = 0.0260, *Thrsp* CHX WT vs CHX S71A p = 0.0043, *Thrsp* WT vs S71A p = 0.0043. Statistical significance is indicated by *p < 0.05 and **p < 0.01, from Mann-Whitney U test. **B)** Motif analyses of promoters of putative PDCD4 target genes (left column) and for comparison, other CHX-insensitive activity-dependent genes (right column) using HOMER software (Heinz et al., 2010). The top panel shows the motif image logos, enrichment, and p-values for the top ten motifs by p-value for activity-dependent upregulated genes, and the bottom panel shows the same but for activity-dependent downregulated genes (only 8 motifs were significant for down-regulated genes).

**Figure S5.**
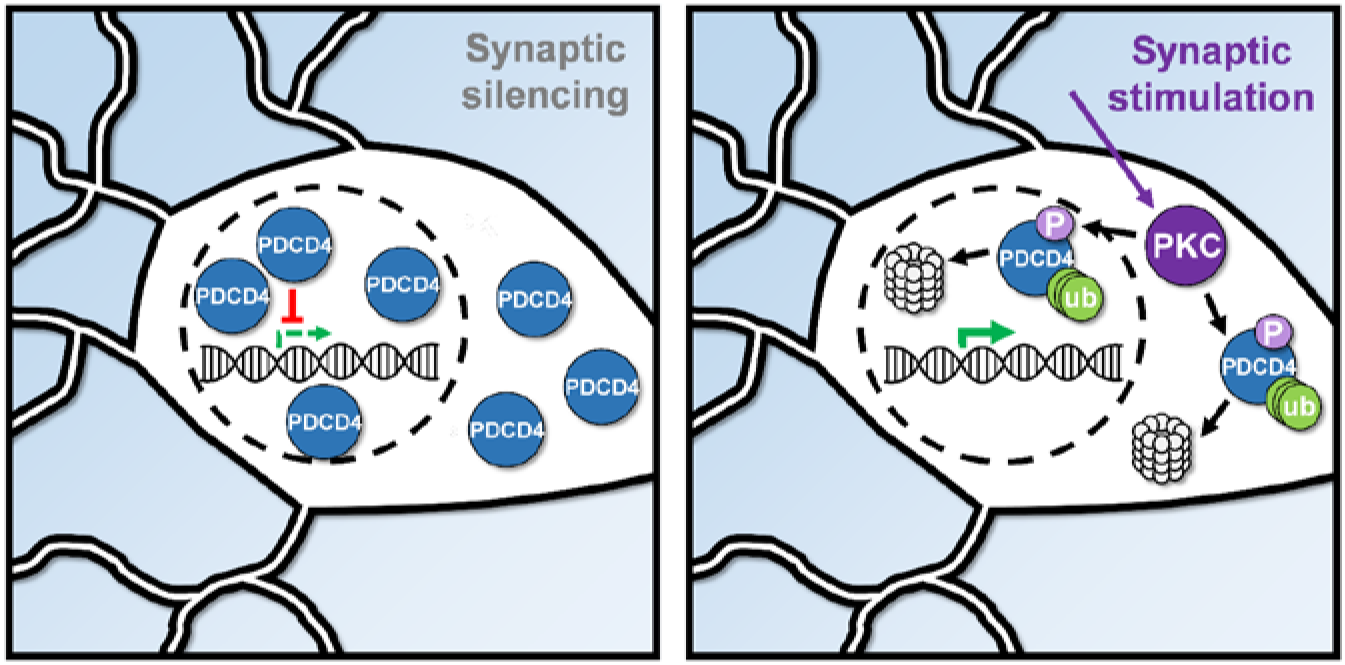
Summary diagram of the activity-dependent proteasome-mediated degradation of PDCD4. In silenced neurons (left), PDCD4 is highly expressed and suppresses the expression of specific genes. In stimulated neurons (right), PDCD4 is phosphorylated by PKC and undergoes proteasome-mediated degradation, thereby facilitating the expression of specific genes important for neuron synaptic function.

## Supplementary Tables

**Table S1. APEX2-NLS Mass Spectrometry of TTX-silenced and Bic-stimulated neurons.**

Mass spectrometry data from the nuclear proteomes of TTX-silenced and Bic-stimulated forebrain cultures, as detected by APEX2 proximity biotinylation. All proteins detected in study (Sheet 1), proteins enriched above the No APEX negative control (Sheet 2), and candidate proteins with differential Bic vs TTX expression (Sheet 3).

**Table S2. Activity-Dependent RNA-Seq of PDCD4 WT- and S71A-transduced neurons.**

RNA-seq data from TTX-silenced and Bic-stimulated forebrain cultures, transduced with PDCD4 WT or S71A, in the presence or absence of CHX. Data for all genes (Sheet 1), data for activity-dependent genes (Sheet 2), and data for putative PDCD4 target genes (Sheet 3).

**Table S3. GO Analysis from PDCD4 RNA-Seq.**

GO analysis data for top 15 terms by FDR for putative PDCD4 target genes (Sheet 1) and other CHX-insensitive activity-dependent genes (Sheet 2).

**Table S4. Primer Sequences Used for Cloning and RT-qPCR.**

